# APP and its intracellular domain modulate Alzheimer’s disease risk gene networks in transgenic *APPsw* and *PSEN1M146I* porcine models

**DOI:** 10.1101/2024.03.21.585176

**Authors:** Mette Habekost, Ebbe T. Poulsen, Jan J. Enghild, Mark Denham, Arne Lund Jørgensen, Per Qvist

**Affiliations:** Department of Biomedicine, Aarhus University; 8000C Aarhus, Denmark; Danish Research Institute of Translational Neuroscience, Nordic EMBL Partnership for Molecular Medicine, Aarhus University; 8000C Aarhus, Denmark; Department of Molecular Biology and Genetics, Aarhus University; 8000C Aarhus, Denmark; Lundbeck Foundation Initiative for Integrative Psychiatric Research, iPSYCH; 8000C Aarhus, Denmark; Centre for Integrative Sequencing, iSEQ, Aarhus University; 8000C Aarhus, Denmark; Centre for Genomics and Personalized Medicine, CGPM, Aarhus University; 8000C Aarhus, Denmark; Developmental and Regenerative Neurobiology, Lund Stem Cell Centre, Department of Experimental Medical Science, Faculty of Medicine, Lund University; 221 84 Lund, Sweden; Center for Translational Neuromedicine, University of Copenhagen Faculty of Health and Medical Sciences; 2200 Copenhagen N, Denmark

## Abstract

Alzheimer’s disease (AD) is a progressive neurodegenerative disorder and the most frequent cause of dementia. The disease has a substantial genetic component comprising both highly penetrant familial mutations (*APP*, *PSEN1,* and *PSEN2*) and sporadic cases with complex genetic etiology. Mutations in *APP* and *PSEN1/2* alter the proteolytic processing of APP to its metabolites, including Aβ and APP Intracellular Domain (AICD). In this study, we use transgenic porcine models carrying the human *APPsw* and *PSEN1M146I* transgenes to demonstrate the pathobiological relevance of transcriptional regulation facilitated by APP and its AICD domain. Through molecular characterization of hippocampal tissue, we describe the differential expression of gene sets that cluster in molecular pathways with translational relevance to AD. We further identify phosphorylated and unphosphorylated AICD in differential complexes with proteins implicated in signal transduction and transcriptional regulation. Integrative genomic analysis of transcriptional changes in somatic cell cultures derived from pigs treated with γ-secretase inhibitor demonstrates the importance of γ-secretase APP processing in transcriptional regulation. Our data supports a model in which APP and, in particular, its AICD domain, modulates gene networks associated with AD pathobiology through interaction with signaling proteins.

**One Sentence Summary:** Utilizing transgenic porcine models, our study reveals that Alzheimer’s disease-related mutations affect neuronal gene expression and highlights the role of the AICD domain of APP in modulating gene networks associated with Alzheimer’s pathobiology.

## INTRODUCTION

Alzheimer’s Disease (AD), the most prevalent form of dementia, represents a significant challenge to global healthcare systems due to its progressive development in the aging population and the absence of curative treatments. Its causes include a combination of genetic, environmental, and lifestyle factors. Mutations in *APP, PSEN1* or *PSEN2* cause fully penetrant familial AD with early-onset (*1–3*), and more than 50 risk loci associated with sporadic AD have been identified in genome-wide association studies (GWASs) (*4*). The familial mutations interfere with the production and processing of APP to amyloid-β (Aβ), a key factor in the formation of neuritic plaques, which is first seen in the hippocampal region of the brain, and is a pathological hallmark of AD. Additionally, several of the risk loci harbor genes involved in the clearance of Aβ (*4, 5*). Such discoveries underscore the central importance of APP’s proteolytic processing into Aβ in the pathogenesis of AD (*6*). While early clinical trials aimed at reducing the Aβ burden in the brain have historically failed to alleviate cognitive symptoms (*7–9*), recent trials using anti-Aβ monoclonal antibodies have lent support to the amyloid hypothesis. Nonetheless, the modest impact of anti-Aβ therapies on disease symptoms raises the question whether other fragments of APP might also contribute to AD pathobiology through mechanisms distinct from those of Aβ (*10*). The type 1 transmembrane glycoprotein, APP, undergoes ectodomain shedding via the action of α- and β-secretases, yielding membrane-bound COOH-terminal fragments (CTFs). Subsequent cleavage by γ-secretase, in turn, produces the non-toxic P3 peptide or Aβ, while releasing the cytoplasmic tail known as APP Intracellular Domain (AICD). AICD has been proposed to function as a transcriptional regulator, potentially in complex with the nuclear adapter protein, FE65 (*11–17*). A range of target genes have been reported, including several involved in APP processing, Aβ clearance, and tau phosphorylation (*13, 18, 19*). Accordingly, AICD is hypothesized to play a key role in disease progression, but the precise molecular mechanism by which AICD may exert regulatory control is a subject of ongoing debate (*11, 20–24*). Understanding such regulatory mechanisms is fundamental for clarifying the pathophysiological cascade of altered APP expression and processing, as well as for the development of effective therapeutic strategies.

Transgenic porcine models expressing mutant forms of human *APP* and *PSEN1* have been developed as a tool for advancing our understanding of AD pathogenesis. Such models offer a more physiologically relevant system than similar rodent models, given the higher anatomical and physiological similarities between the porcine and the human brain. Notably, the human and porcine brains share identical secretase cleavage sites in APP and both harbor the 3R tau isoform, which is absent in the rodent brain (*25*). Here, we demonstrate in transgenic porcine models expressing the *APPsw* and *PSEN1M146I* mutations (*26–29*) the molecular impact and pathobiological relevance of transcriptional regulation associated with APP processing, including the release of its AICD domain.

## RESULTS

### Molecular profiling reveals shared effects in transgenic pig hippocampi

To assess the molecular impact of altered APP production and processing, we performed whole transcriptome profiling in macrodissected hippocampal tissue samples from wild-type (WT) pigs and transgenic pigs that express one copy of human *PSEN1* with the M146I mutation (*PSEN1M146I*; hereafter simply PS1) or human *APP* with the Swedish double-mutation K670N/M671L (*APPsw*) in combination with *PSEN1M146I* (hereafter simply APP/PS1) (**Fig. 1A and Table S1, S2**) (*15, 27, 30*). By comparing mRNA levels between PS1 and WT, and APP/PS1 and WT samples, we identified, respectively, 1,539 nominally differentially expressed genes (DEGs_Pnom_, P < 0.05) and 599 DEGs_Pnom_ (P < 0.05) – of which correspondingly, 65 and 61 were significant after correction for multiple testing (DEGs_Padj_, P < 0.05) (**Fig. 1B**). A significant overlap of DEGs_Padj_ were identified in PS1 and APP/PS1 pigs (14 shared; **Fig. 1B**; **Table S1, S2**; Fisher Exact pathway test P < 0.05) and expression characteristics of the combined list of DEGs_Padj_ across samples suggest a shared molecular effect of transgene expression in hippocampal samples derived from the two groups of transgenic pigs (**Fig. 1C; Fig. S1A)**. Since the transgenic pigs carry human APP and PS1 transgenes, the alignment of RNA sequencing reads to the porcine genome did not effectively detect the differential expression of these genes. Furthermore, the expression level of cell-type-specific genes *OLIG1* (oligodendrocytes), *GFAP* (astrocytes) and *RBFOX3* (neurons) showed no significant variance across the examined samples (**Fig. S1B, C**). Additionally, analysis of DEGs_Padj_ with cell-type-specific expression (**Fig. 1C**) (*31*) suggested that the changes in gene expression were not attributable to differences in cellular composition among the tissue samples. Of the more than 50 risk loci reaching genome-wide significance for association with AD (*4*), 35 were expressed in the hippocampal tissues studied, but none of the risk genes were found to be differentially expressed after correction for multiple testing. A gene-set association analysis utilizing summary statistics from the largest-to-date AD GWAS (*32*), highlights the etiopathological relevance of the identified transcriptomic changes, as particularly the PS1 DEGs_Padj_ set displays a significant association (P = 0.03). In addition, several risk genes (*NDUFAF7, ADAM10, APP, DSG2, CLU, SORL1, BIN1, MAPT, PICALM, PPARGC1A* and *PLCG2*) were identified to have known experimental or database interactions with members of the DEGs_Padj_ list in networks linked to mitochondrial oxidative metabolism, G protein signaling, microtubule formation, and lipid metabolism (**Fig. S2**). Adding to this notion, functional analysis of DEGs_Pnom_ genesets using Ingenuity Pathway Analysis (IPA) tool revealed enrichment of pathways related to signaling transduction, oxidative phosphorylation and mitochondrial functions (**Fig. 1D**). Notably, the fold changes of genes annotated to the ‘Oxidative Phosphorylation’ gene set predicted an activation of this pathway in the hippocampus of APP/PS1 pigs (Z-score = 2.121) (**Fig. 1D** and **Table S3, S4**). The highest ranking pathways in PS1 pigs with significant activation scores were ‘PI3K Signaling in B Lymphocytes’ (Z-score = 2.449), ‘RhoGDI Signaling’ (Z-score = 2.449), and ‘G beta Gamma Signaling’ (Z-score = 2), while ‘Signaling by Rho Family GTPases’ was predicted to be inhibited (Z-score = -2.646) (**Fig. 1D** and **Table S3, S4)**.

**Fig. 1.**
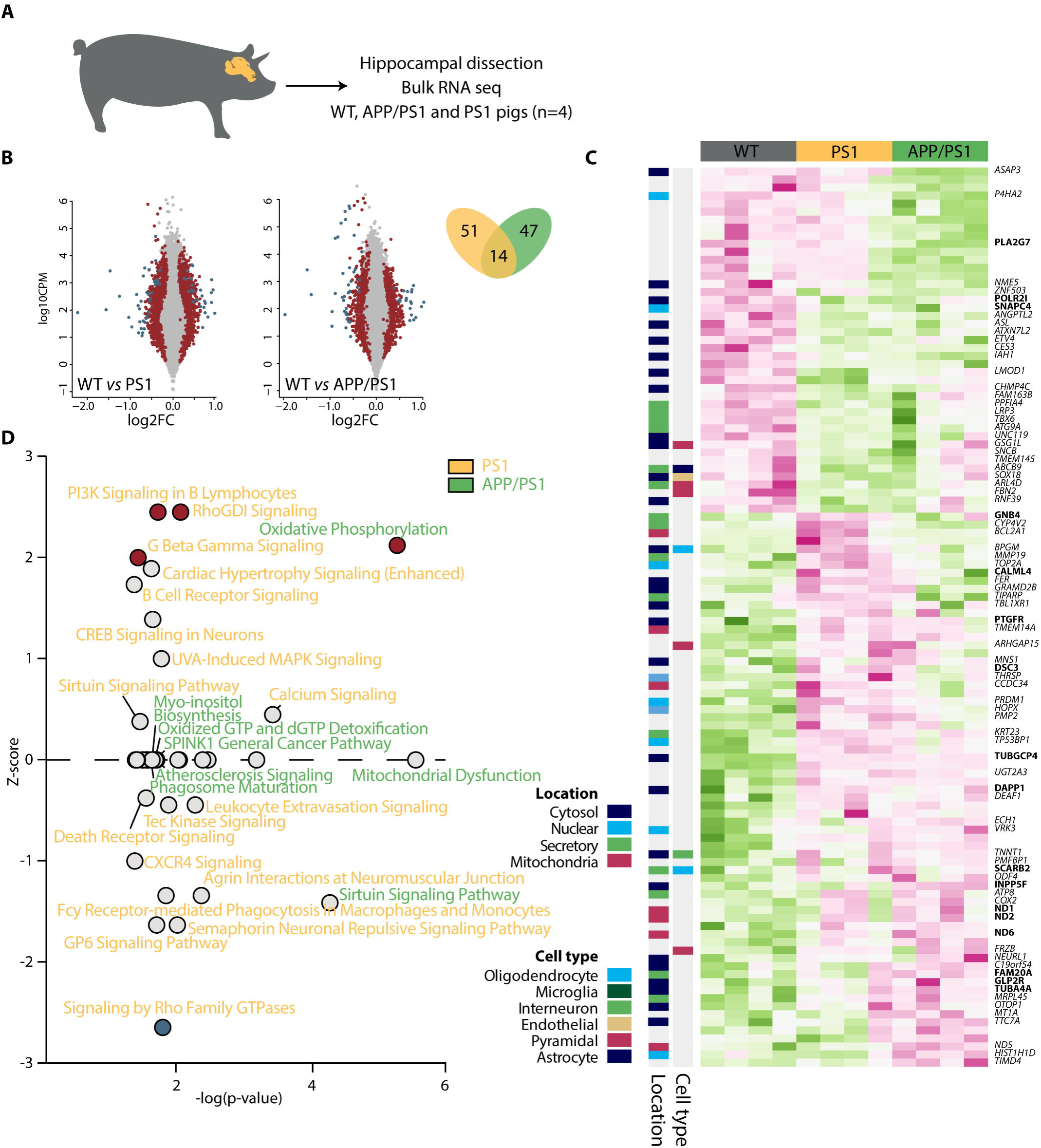
Shared molecular effects in transgenic hippocampal tissues. **(A)** Schematic overview. **(B)** Log2 fold change (FC) in x-axis and log2 counts per million (CPM) in y-axis after deseq2 differential expression analysis of PS1 (yellow) and APP/PS1 (green) against WT. P nominal < 0.05 in red, P adjusted < 0.05 in blue. Venn diagram of differentially expressed genes in PS1 (yellow) and APP/PS1 (green). **(C)** Heatmap of log2 transformed and mean scaled TPM expression values of the combined list of DEG_Padj_ (PS1 vs WT and APP/PS1 vs WT) across WT, PS1 and APP/PS1 hippocampal tissues. Subcellular location and cell type-specific expression analysis of DEGs_Padj_ listed as color codes on the left. Only annotated gene symbols are shown. Gene symbols highlighted in bold denote genes that have either experimental or database interactions with known risk loci as detailed in Fig. S2. **(D)** Enriched pathways plotted as -log(pvalue) in x-axis and Z-score in y-axis after Ingenuity Pathway Analysis (IPA) of DEGs identified in PS1 (yellow) or APP/PS1 (green) pigs. Red or blue dots indicate the significant Z-scores above 2 or below -2, respectively.

### Proteomic analysis identifies AICD binding partners in the hippocampus

The _682_YENPTY_687_ (note APP695 numbering) motif within the AICD domain of APP plays a crucial role in protein-protein interactions (*29*). Particularly, phosphorylation of the tyrosine residue at position 682 (Y_682_) influences the binding affinity and specificity for various intracellular proteins, including those with Src homology 2 (SH2) domains and Phosphotyrosine-binding (PTB) domains. To explore the putative role of AICD in gene regulation associated with PS1 and APP/PS1 in the hippocampus, we utilized a pull-down assay employing synthetic peptides corresponding to the YENPTY motif to compare the interactomes of phosphorylated (AICD-pY) and non-phosphorylated AICD (AICD-Y) at Y_682_ (**Fig. 2A**). Negative controls included peptides with Y_682_ substituted with glycine (AICD-YG), and a scramble amino acid sequence (SCR). The peptides were incubated with WT, PS1, and APP/PS1 pig hippocampal lysates, and the interaction partners were subsequently identified by LC-MS/MS analysis (**Fig. 2B**).

**Fig. 2.**
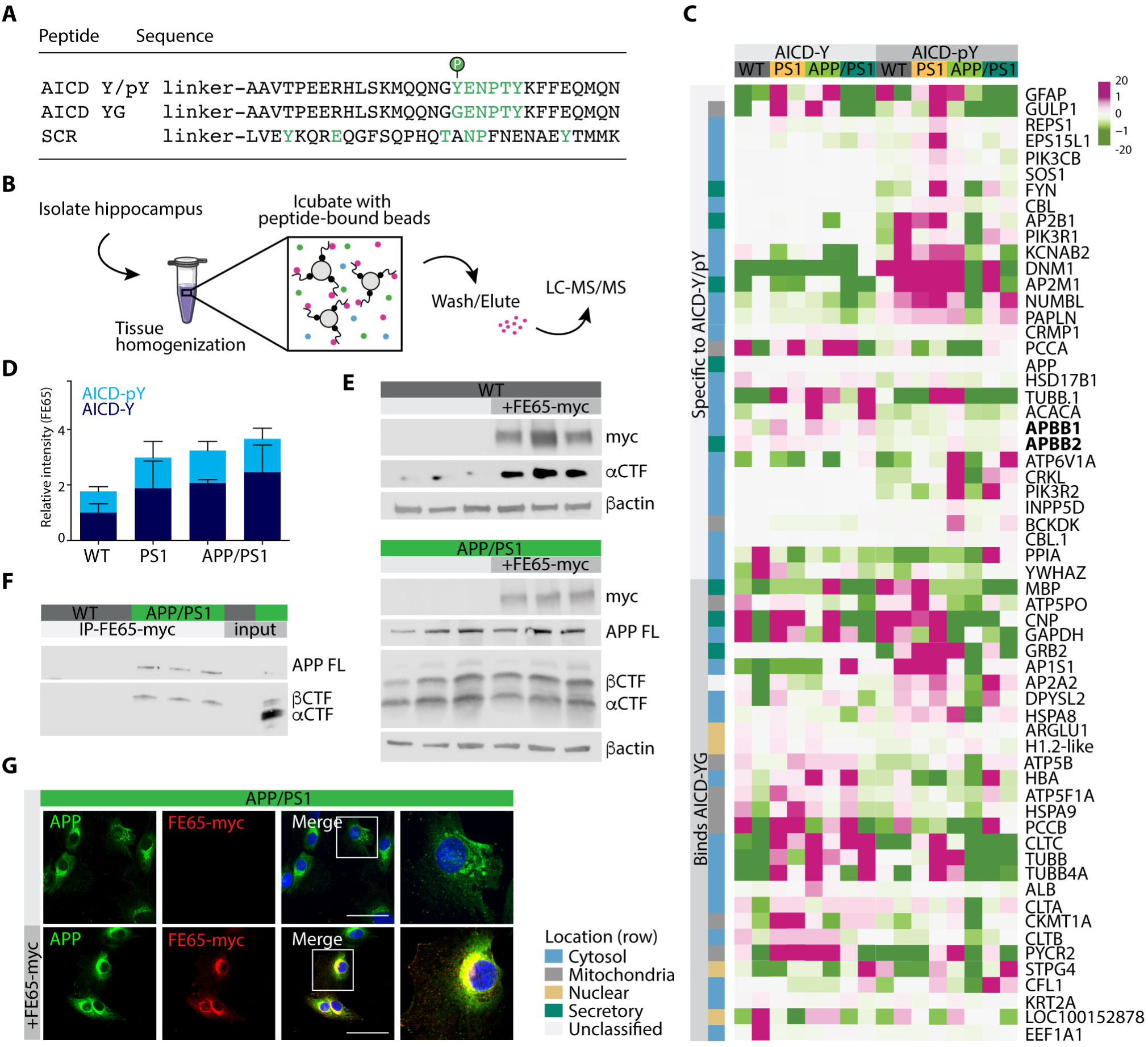
Identification of AICD binding partners in hippocampal tissue. **(A)** Synthetic peptides used for protein pull-down experiments. **(B)** Schematic illustration of experimental flow. **(C)** Proteins identified by LC-MS/MS of AICD-mimicking synthetic peptide pull-down experiments in hippocampal tissues from WT, PS1 and APP/PS1 transgenic pigs. Proteins plotted as heatmap without row normalization. Color intensity (green to pink) indicates emPAI values. Proteins included did not bind to SCR peptides and were identified in all genotypes. Subcellular location analysis of the identified proteins listed as color codes on the left. APBB1 (FE65) and APBB2 (FE65L1) indicated in bold. **(D)** Targeted MS method monitoring FE65 to measure relative quantities of the protein in the peptide pull-down experiment. **(E)** Western blot analysis of total lysates from WT and APP/PS1 porcine fibroblasts overexpressing myc-tagged human FE65 using anti-APP-c-terminal antibody (clone C1/6.1), anti-myc (clone 9E10) and βactin as loading control. *n*=3 biological replicates. **(F)** Co-immunoprecipitation analysis of fibroblasts from WT (grey) and APP/PS1 (green) fibroblasts overexpressing myc-tagged human FE65. Immunoprecipitations were performed with anti-myc antibodies and analyzed with anti-APP-c-terminal antibody. *n*=3 biological replicates. **(G)** Confocal microscopy analysis of APP/PS1 fibroblasts overexpressing myc-tagged human FE65 using anti-myc antibodies (in red) and anti-APP-c-terminal antibody (in green). Scale bar, 50μm. APP FL, APP full length, CTF, C-terminal Fragment.

We identified 60 proteins consistently pulled down in all samples but not in SCR controls (**Fig. 2C, Table S5**). More than half of these interactions were dependent on Y_682,_ as they did not bind to the AICD-YG peptide. Notable APP interaction partners such as 14-3-3 (YWHAZ), GULP1, SOS1, FYN, NUMBL, FE65 (APBB1), FE65L1 (APBB2), GRB2, SHIP (INPP5D) (*33*), were among the identified proteins. A subset of these preferentially bound to phosphorylated (e.g. NUMBL, FYN, SHIP, SOS1) or unphosphorylated (e.g. FE65, FE65L1, GULP1) AICD peptides (**Fig. 2C**).

Subcellular location analysis using the SubCellBarCode resource (*34*) revealed that the majority of interaction partners are predicted to localize in the cytosol and mitochondria (**Fig. 2C**). However, supporting a role for AICD in transcriptional regulation, some have predicted nuclear actions (**Fig. 2C**). To provide validation, we performed LC-parallel reaction monitoring (PRM)-MS, with a focus on FE65. This analysis confirmed the identity of the protein and demonstrated the tendency to preferably bind AICD-Y over AICD-pY peptides (**Fig. 2D**). To further validate the LC-MS/MS analysis, co-immunoprecipitation was performed in FE65-myc transfected WT and APP/PS1 fibroblasts. In WT fibroblasts, expression of *FE65*-*myc* increased APP-CTF protein levels, while APP processing remained unchanged in APP/PS1 fibroblasts (**Fig. 2E)**. Immunoprecipitation using myc-tagged beads followed by western blotting pulled down APP in APP/PS1 fibroblasts, while APP was below detection level in WT cells (**Fig. 2F**). We specifically detected the APP β-CTF, which is derived from the initial cleavage step in the amyloidogenic processing of APP by β-secretase. Co-localization analysis confirmed co-occurrence of APP and myc-tagged FE65 in the cytosol **(Fig. 2G**), confirming the efficacy of our pull-down assay in identifying proteins interacting with AICD.

### Transgene expression influences AD-related pathways in fibroblasts

To explore and modulate the transcriptomic impact of *APPsw* and *PSEN1M146I* transgene expression and APP processing, fibroblasts were isolated from WT and APP/PS1 pigs and subjected to transcriptomic profiling (**Fig. 3A**). As APP overexpression is known to regulate cell proliferation (*35*), we limited our analysis to genes with translational relevance to postmitotic neurons, by excluding raw counts not mapping to neuron-expressed genes. Using this approach, we identified 1460 DEGs_Pnom,_ of which 502 were significant following correction for multiple testing (DEGs_Padj_; **Fig. 3B, Table S6**). Despite filtering, we detected enrichment of pathways related to cell proliferation (**Fig. 3C**), including ‘Cell Cycle Control of Chromosomal Replication’ (-log(P) = 10.9, Z score = -3.2), but we also detected enrichment of pathways with direct relevance to AD pathobiology and signal transduction including ‘Neuroinflammation Signaling Pathway’ (-log(P) = 1.48, Z score = 1.732), ‘Amyloid Processing’ (-log(P) = 2.33), ‘14-3-3-mediated Signaling’ (-log(P) = 1.21), ‘RAN Signaling’(-log(P) = 3.36, Z score = -2) and ‘P53 Signaling’ (-log(P) = 2.84, Z score = 0.45) amongst others (**Fig. 3C, Table S7**).

**Fig. 3.**
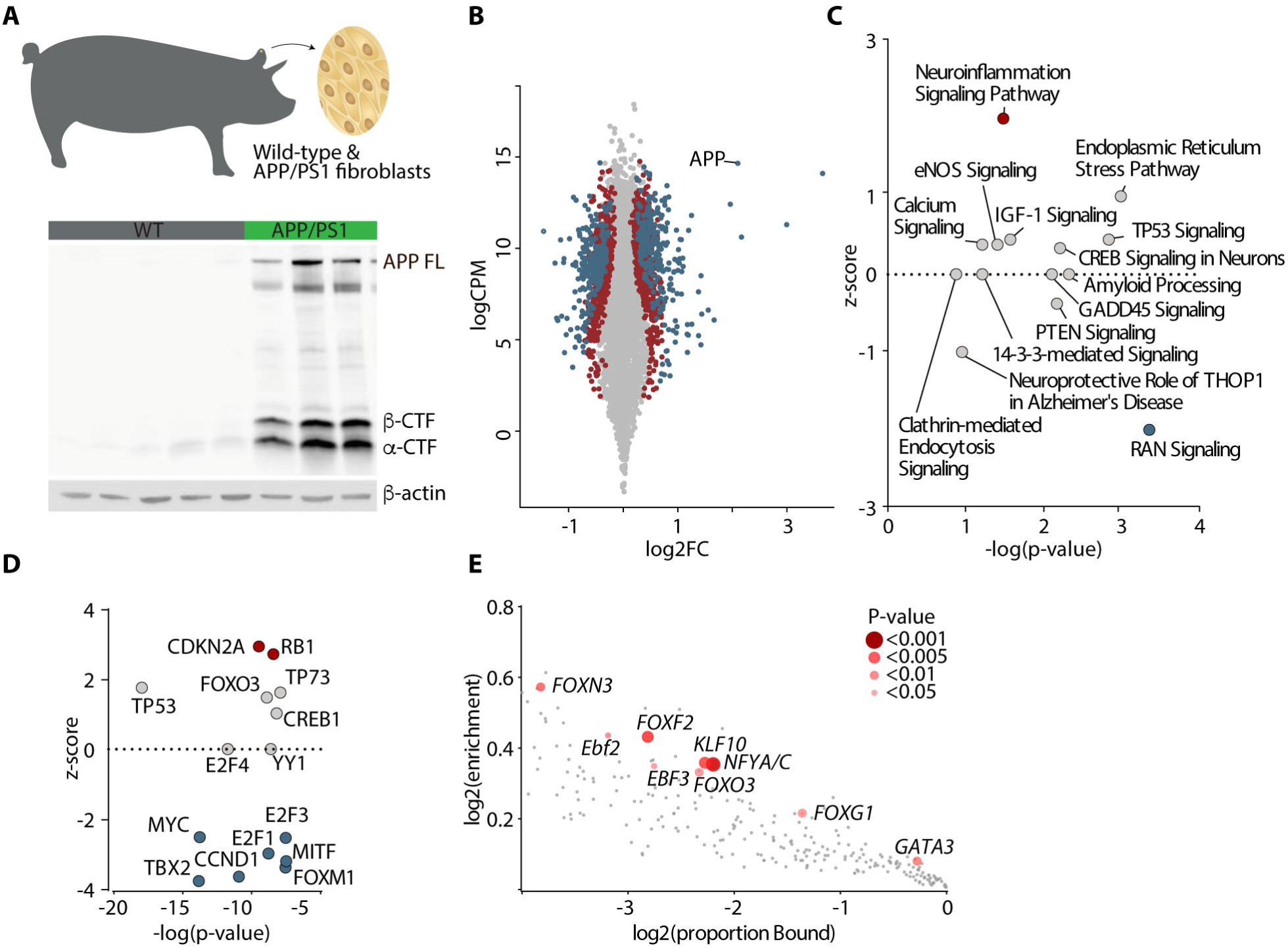
APP/PS1 expression in fibroblasts modulates AD-related pathways. **(A)** Western blot analysis of total lysates from WT and APP/PS1 porcine fibroblasts using anti-APP-c-terminal antibody (clone C1/6.1). βactin as loading control. *n*=3-5 biological replicates. **(B)** Log2 fold change (FC) in x-axis and log2 counts per million (CPM) in y-axis after deseq2 differential expression analysis of APP/PS1 against WT fibroblasts. P nominal < 0.05 in red, P adjusted < 0.05 in blue. **(C)** Enriched pathways plotted as -log(pvalue) in x-axis and Z-score in y-axis after Ingenuity Pathway Analysis (IPA) of DEGs identified in APP/PS1 fibroblasts. **(D)** Upstream Regulators plotted as -log(pvalue) in x-axis and Z-score in y-axis after Ingenuity Pathway Analysis (IPA) of DEGs identified in APP/PS1 fibroblasts. Red or blue dots indicate the significant Z-scores above 2 or below -2, respectively. **(E)** Ciiider analysis identification of transcription factor binding sites within the promotor sequences (1500bp upstream and 500bp downstream of TSS (Homo sapiens GRCh38.94)) in the list of DEG_Padj._

The IPA upstream transcription regulator analysis of DEGs_Padj_ identified several transcription factors potentially driving the observed gene expression changes. Among these, FE65 emerged as a notable upstream regulator (APBB1; P = 0.044)(**Table S8**). Additionally, several other transcription factors known to either interact with AICD or be influenced by it, such as TP53, TP73, MYC and FOXO3, were identified in this analysis (**Fig. 3D**). The significance of genes regulated by FOX group transcription factors, particularly FOXO3, was further supported by a transcription factor binding site (TFBS) enrichment analysis conducted on DEG_Padj_ promoter sequences (**Fig. 3E, Table S9**). Notably, hierarchical clustering of expression data for genes with FOXO3 TFBS in their promotor regions distinctively segregated WT and APP/PS1 fibroblasts samples in two separate clusters (**Fig. S3**).

### Secretase-inhibition alters gene expression in transgenic pig fibroblasts

Building on the findings from the upstream transcription factor analysis, we considered the postulated role of liberated AICD as a transcription factor, particularly when in complex with the nuclear adapter protein, FE65 (*11, 20*). It has been reported that AICD generated from APP-CTFs through the action of γ−secretase predominantly originates from β−secretase produced CTFs (*21, 36, 37*). We therefore examined the transcriptional impacts of both β− and γ−secretase inhibition in WT and APP/PS1 fibroblasts (**Fig. 4A**). In APP/PS1 γ-secretase inhibited cells, we identified a total of 231 DEG_Padj_ compared to their non-inhibited controls (**Fig. 4B, C, Table S10)**, while only *LDLR* was identified as a DEG_Padj_ in APP/PS1 β-secretase inhibited cells (**Fig. 4B, C**). No DEG_Padj_ were identified in WT cells treated with either inhibitor (**Fig. 4B, C**). There was a larger overlap of DEG_Padj_ induced by APP/PS1 transgene expression and γ-secretase inhibition than expected by chance (**Fig. 4D**; Fisher Exact test, P < 0.05) and γ-secretase inhibition partially normalized APP/PS1 associated gene expression changes (**Fig. S4**). Interestingly, genes impacted by γ-secretase inhibition in APP/PS1 samples displayed a very significant gene set association to AD (P < 0.001), but much weaker or no association to other brain disorders tested (Parkinson’s disease (P = 0.06) (*38*) Tourette’s syndrome (P = 0.44) (*39*) and broad sense psychiatric illness captured in a recent cross disorder GWAS (P = 0.67) (*40*)), making it unlikely that the association is attributed to the neuron-expressed gene-enriched nature of the gene set. Corroborating the observation that impeding AICD release can counteract the transcriptional effects induced by APP/PS1 transgene expression, our upstream regulator analysis once again highlighted FOXO3 (P = 3.27E-06, **Fig. 3E, Table S11**) as a primary regulator influencing gene expression changes associated with γ-secretase inhibition. Hierarchical clustering of FOXO3 target genes further supported this finding, as it distinctly segregated γ-secretase inhibitor-treated and untreated APP/PS1-derived fibroblasts into different clusters. This, in turn, was not evident in WT fibroblasts, which suggests that a specific transcriptional response to γ-secretase inhibition in the APP/PS1 fibroblasts was mediated through FOXO3 (**Fig. S5**). Importantly, the expression level of FOXO3 was not affected by γ-secretase inhibition (P = 0.89), suggesting that FOXO3 activity, rather than expression level, is responsible for the observed effect.

**Fig. 4.**
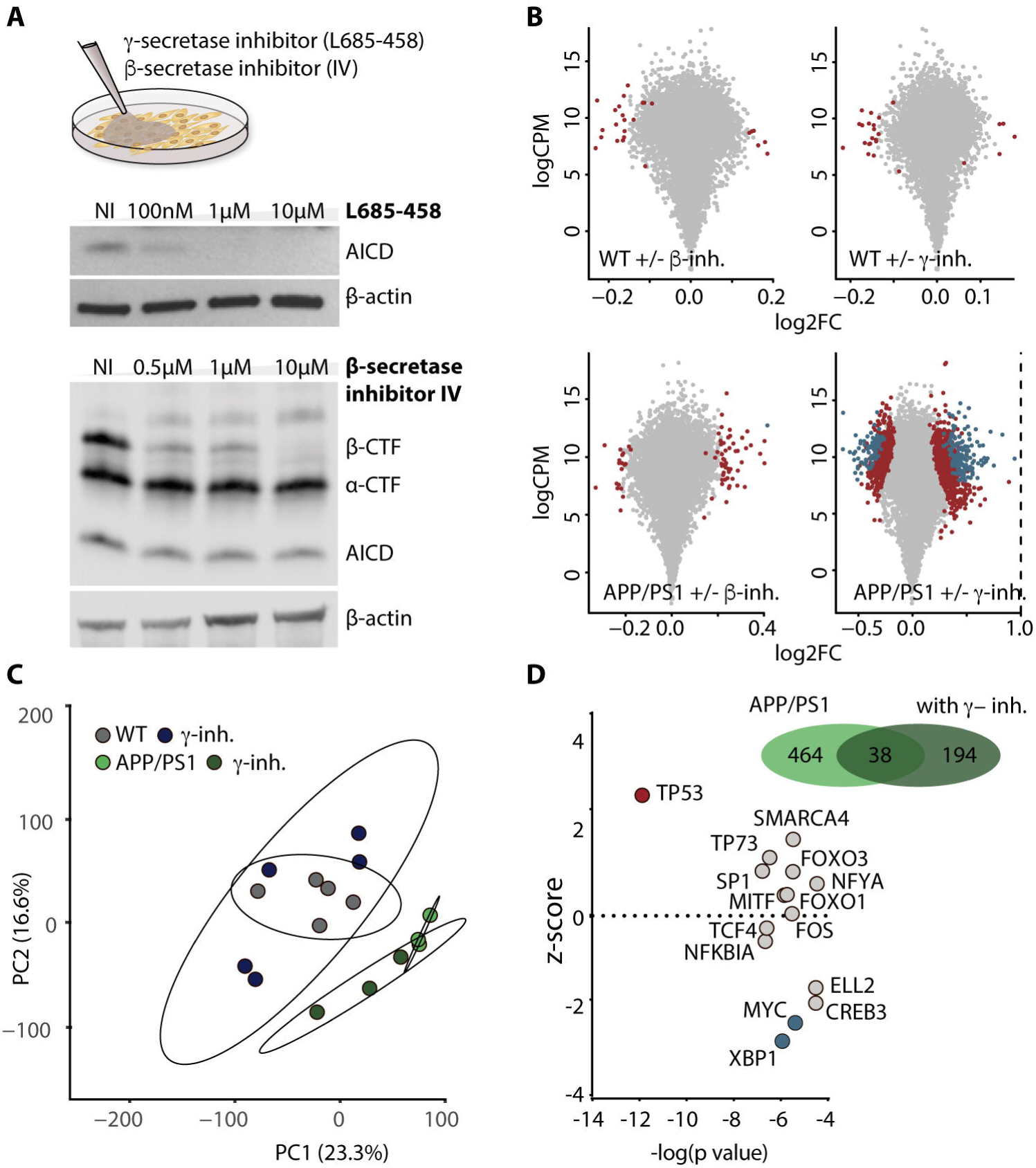
γ−secretase inhibition alters gene expression in transgenic fibroblasts. **(A)** Western blot analysis of total lysates from APP/PS1 porcine fibroblasts treated with increasing concentrations of γ− (L685-458; 100nM-10μM) and β−secretase (0.5−10 μM) pharmacological inhibitors using anti-APP-c-terminal antibody (clone C1/6.1). βactin as loading control. **(B)** Log2 fold change (FC) in x-axis and log2 counts per million (CPM) in y-axis after deseq2 differential expression analysis of APP/PS1 and WT fibroblasts treated with γ− and β−secretase inhibitors. P nominal < 0.05 in red, P adjusted < 0.05 in blue. **(C)** Whole transcriptome PCA plot of APP/PS1 and WT fibroblasts treated with γ−secretase inhibitor. **(D)** Venn diagram of differentially expressed genes in APP/PS1 (green) and APP/PS1 γ−inhibited fibroblasts (dark green), and upstream regulators plotted as -log(p-value) in x-axis and z-score in y-axis after Ingenuity Pathway Analysis (IPA) of DEGs identified in APP/PS1 fibroblasts treated with γ−secretase inhibitor. Red or blue dots indicate the significant Z-scores above 2 or below -2, respectively.

Among previously reported AICD target genes (*HES1, TAGLN, TPM1, GSK3B, LRP1, EGFR, CCND1, CCNB1, DDIT3, SLC17A6, TRPC5, PTCH1, SPTLC2, CLU, MAT2A, MYC, STMN1, PML, PPARGC1A, GADD45G, TTR, AGPS, TP53, TP73*) (*33*), we noted alterations in the expression levels of *LRP1* (P = 0.02), *CLU* (P = 0.001) and *SPTLC2* (P = 0.04) under conditions where γ-secretase inhibition impedes AICD release (**Table S10**). However, it is critical to consider that APP/PS1 fibroblasts have elevated AICD levels in contrast to the nominal levels in WT cells. Despite this variance in AICD abundance, no significant changes were observed in the expression of *LRP1*, *CLU,* and *SPTLC2* between APP/PS1 and WT fibroblasts (**Table S6**). This suggests that AICD may not directly function as a transcription factor in regulating these specific genes.

### Protein network analysis links AICD to FOXO3 regulation

To understand how AICD’s interactions may drive the transcriptional changes observed in our models, we conducted a protein-protein interaction network analysis between putative upstream regulators (in blue; **Fig. 5A**) and AICD’s identified interaction proteins (in yellow; **Fig. 5A**). The network revealed strong direct as well as indirect connections between APP and upstream regulators, suggesting their potential role in facilitating APP/PS1-induced transcriptional changes. Given the observed alteration in FOXO3 expression levels in APP/PS1 fibroblasts relative to WT controls (Padj = 0.008) and its deregulated activity upon γ-secretase inhibition, we further explored the relationship between FOXO3 and APP. The phosphorylation state of FOXO3 is crucial in regulating its transcriptional activity, which is modulated by several pathways including the phosphatidylinositol 3-kinase (PI3K) and protein kinase B (AKT) signaling cascade. PI3K activates AKT (phosphorylated at threonine 308 and serine 473), leading to the phosphorylation of FOXO3 and subsequent inhibition of its nuclear localization (*41*). Intriguingly, our study identified PIK3R1 (P85) and PIK3CB (P110) as phosphorylation-dependent binding partners of AICD (**Fig. 2C** and **Fig. 5B**). To assess the impact on the AKT signaling pathway, we conducted western blot analyses on APP/PS1 and WT fibroblasts, utilizing antibodies specific for phospho-AKT (Ser473) and total AKT. Our results revealed a reduction in phospho-AKT protein levels in APP/PS1 fibroblasts compared to WT. This finding was substantiated by treating the fibroblasts with LY294992, a selective inhibitor of PI3K-dependent AKT phosphorylation (**Fig. 5C and D**) (*42*). These observations suggest the PI3K/AKT pathway as a potential intermediary between APP and the transcriptional alterations mediated by FOXO3.

**Fig. 5.**
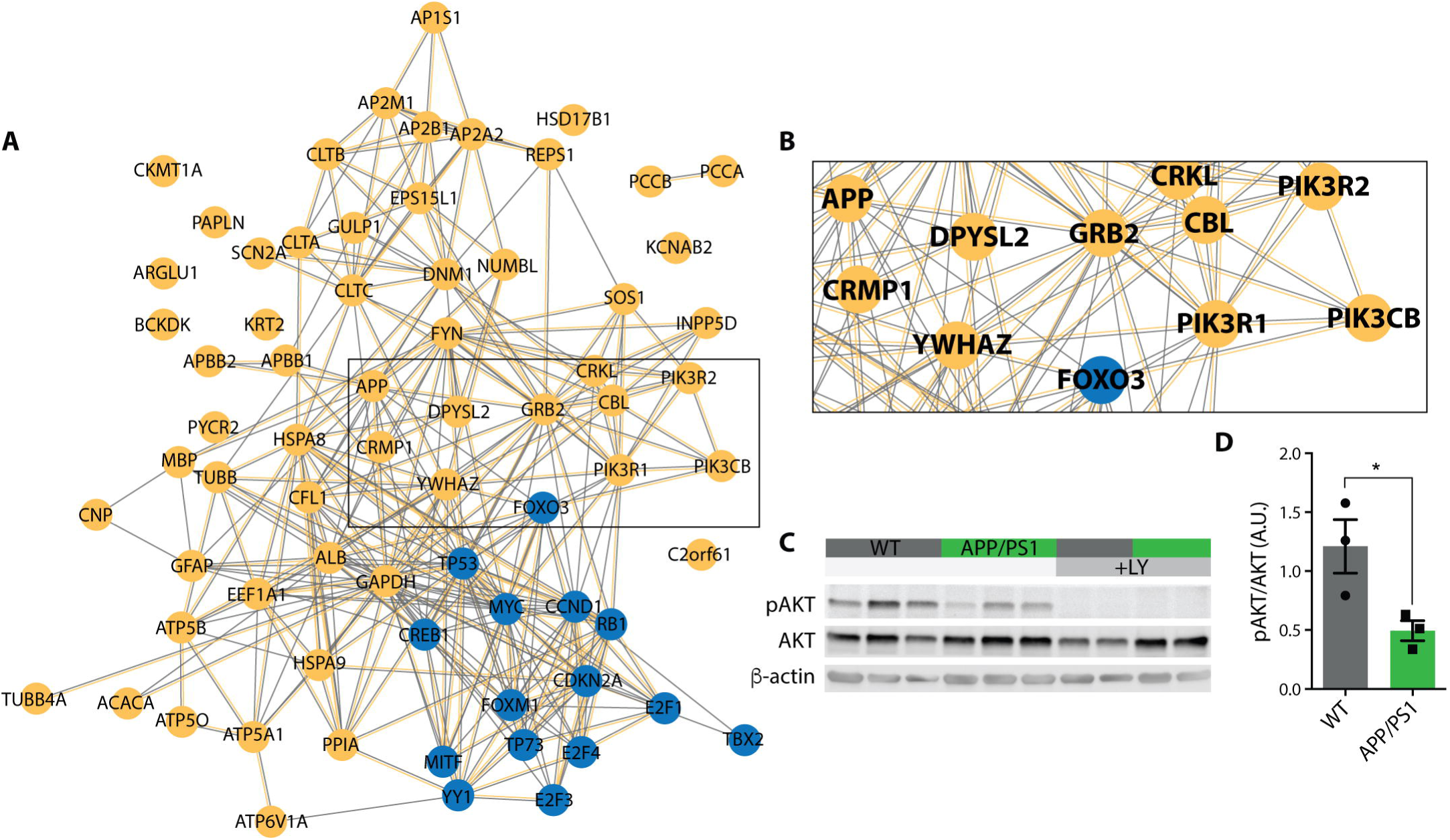
Protein-protein interaction network points to altered PI3K/AKT phosphorylation in APP/PS1 fibroblasts. **(A)** STRING protein-protein interaction network of APP interactome (in yellow) and potential upstream regulators (in blue) identified in Fig. 2C. Interaction sources: database (grey) and experiments (yellow). **(B)** Enlarged image of the marked area in A. **(C)** Western blot analysis of total lysates from WT (grey) and APP/PS1 (green) porcine fibroblasts with or without treatment with LY294002 (50μM) using anti-phospho AKT (Ser473) (clone D9E) and anti-AKT antibodies. βactin as loading control. *n*=3 or 2 biological replicates. **(D)** Quantification of AKT phosphorylation to total AKT levels in C.

## DISCUSSION

In this study, we assess the molecular impact of transgenic *APPsw* and *PSEN1M146I* overexpression in a porcine model with translational relevance to the neuropathology of AD. We observe widespread changes in hippocampal gene expression in transgenic pigs, that display gene set association to AD and cluster in molecular pathways that align with known AD pathobiology (*43–45*). The activation of the oxidative phosphorylation pathway resonates with the growing body of evidence suggesting mitochondrial dysfunction as a key player in AD pathology (*46*). Importantly, these changes in mRNA levels do not appear to result from differences in cellular composition or changes in neuronal activity, as evidenced by the shared transcriptomic signatures in hippocampal tissue and fibroblasts derived from transgenic APP/PS1 pigs. Particularly, our characterization of *APPsw* and *PSEN1M146I* induced gene expression changes across tissue and cells consistently points to altered aging related signaling pathways such as mitochondrial dysfunction, FOXO3 transcription factor, sirtuin and P53 signaling.

The intracellular domain of APP has garnered attention for its role in signaling and gene regulation. Our study reveals that endogenous FE65, a protein with a putative role in AICD-mediated transcriptional regulation, binds to AICD in hippocampal samples. This interaction has previously been difficult to detect in proteomic studies without overexpression of FE65 (*47*). Contrary to conflicting reports on the impact of Y_682_ phosphorylation (*48, 49*), our results align with studies indicating that FE65 can bind to AICD irrespective of phosphorylation status. This also aligns with previous reports suggesting that posttranslational modifications on T_668_ upstream YENPTY are the main regulators of the interaction (*50–53*). In our experimental systems, we observed that FE65 associated with the β-CTFs in the cytosol, suggesting a preference for interactions with APP fragments generated by β-secretase cleavage. While FE65 emerged as a potential upstream regulator in our transcriptomic analysis of cells from APP/PS1 pigs, our data did not support a direct function as a transcriptional regulator for liberated AICD in complex with FE65. Hence, further experiments are needed to elucidate the complexity of AICD’s interactions with FE65.

Instead, our findings propose that AICD influences gene transcription indirectly through association with protein networks that mediate downstream transcriptional activity. Particularly, FOXO3 emerges as a probable regulator of a specific group of DEGs, as supported in our study by multiple analytical approaches. Notably, inhibiting γ-secretase dependent release of AICD in APP/PS1 fibroblasts appears to partially normalize the expression of genes targeted by FOXO3. Interestingly, FOXO3 has been highlighted in a range of other studies as a transcription factor modulated by AICD (*54, 55*). Direct interactions between FOXO3 and AICD have been reported (*56*), but we did not detect such complexes in our hippocampal tissue samples. Our findings suggest a more indirect pathway through which AICD exerts influence over FOXO3 activity.

The activity of FOXO3 is predominantly controlled by phosphorylation, which restricts its nuclear entry and blocks the transcription of pro-apoptotic genes (*57, 58*). AKT is a kinase known to inhibit FOXO3 and is part of the PI3K/AKT signaling pathway. This pathway is initiated by PI3K, which consists of P110 catalytic and P85 regulatory subunits (*41, 59*). We observed in hippocampal samples that these subunits form a complex with phosphorylated AICD and identified a corresponding decrease in AKT phosphorylation in APP/PS1 fibroblasts. Previous reports have similarly coupled AICD to increased FOXO3 activity following the loss of phosphorylated AKT (*60*). Diminished PI3K/AKT pathway activity is also associated with the upregulation of GSK3β, a kinase implicated in tau phosphorylation and Aβ accumulation (*61*). Thus, AICD, through its interactions, appears to influence molecular networks related to AD pathogenesis. However, the specifics of how AICD’s association with the PI3K complex leads to a decrease in AKT activation remains undefined. Future investigations are necessary to elucidate the mechanisms and functional implications of AICD’s phosphorylation-dependent interactions and the impact of γ-secretase processing on these networks.

## MATERIALS AND METHODS

### Study design

The study aimed to investigate the role of transcriptional regulation facilitated by APP and its AICD domain in the pathobiology of Alzheimer’s disease. We obtained samples from single- and double-transgenic Göttingen minipigs carrying three (genotype APP/PS1#592) or more (genotypes APP/PS1#590 and #509) copies of human APP695 cDNA with the Lys670Asn/Met671Leu (*APPsw*) double-mutation and/or one copy of human PSEN1 cDNA with the Met146Ile (genotype PS1) mutation. Detailed genotypic descriptions are previously reported (*26, 27*). The samples utilized in our study are outlined in **Table 1**. Inclusion criteria for samples were based on genotype and age. There were both male and female samples across all genotypes.

**Table 1.** Overview of samples used in this study. . RNAseq, RNA sequencing; MS, Mass Spectrometry; P, Passage number.

| Genotype | ID and gender | Sample | Age | Protocol |
| --- | --- | --- | --- | --- |
| WT | 1E ♀ | Hippocampus | 3 years 6 months | RNAseq |
| WT | 2E ♀ | Hippocampus | 3 years 6 months | RNAseq |
| WT | 317245 ♂ | Hippocampus | 3 years | RNAseq/MS |
| WT | 852 ♀ | Hippocampus | 3 years 2 months | RNAseq |
| WT | 498 ♂ | Fibroblast/<br>Hippocampus | 5 months; P5 | RNAseq/MS |
| WT | 499 ♀ | Fibroblast | 5 months; P5 | RNAseq |
| WT | 500 ♀ | Fibroblast | 5 months; P5 | RNAseq |
| WT (A) | 522 ♂ | Fibroblast | 6 months; P9 | RNAseq |
| WT (A) | 512 ♂ | Fibroblast | 10 months; P9 | RNAseq |
| PS1 | 508 ♂ | Hippocampus | 3 years 1 months | RNAseq |
| PS1 | 538 ♂ | Hippocampus | 2 years 1 months | RNAseq |
| PS1 | 540 ♂ | Hippocampus | 3 years 8 months | RNAseq |
| PS1 | 495 ♀ | Hippocampus | 5 years 1 months | RNAseq |
| PS1 | 507 ♂ | Hippocampus | 11 months | MS |
| PS1 | 503 ♂ | Hippocampus | 3 months | MS |
| APP/PS1 (592) | 654 ♀ | Hippocampus | 10 months | MS |
| APP/PS1 (592) | 577 ♀ | Hippocampus | Newborn | MS |
| APP/PS1 (591) | 6027 ♀ | Hippocampus | 3 years 6 months | RNAseq |
| APP/PS1 (592) | 6170 ♀ | Hippocampus | 3 years 7 months | RNAseq |
| APP/PS1 (592) | 6172 ♀ | Hippocampus | 4 years 8 months | RNAseq |
| APP/PS1 (592) | 6171 ♀ | Hippocampus | 5 years 4 months | RNAseq |
| APP/PS1 (590) | 590 ♀ | Hippocampus | Newborn | MS |
| APP/PS1 (590) | 606 ♀ | Fibroblast | Newborn; P5 | RNAseq |
| APP/PS1 (590) | 607 ♀ | Fibroblast | Newborn; P5 | RNAseq |
| APP/PS1 (509) | 560 ♂ | Hippocampus | Newborn | MS |
| APP/PS1 (509) | 550 ♂ | Fibroblast | Newborn; P5 | RNAseq |
| APP/PS1 (509) | 557 ♂ | Fibroblast | Newborn; P5 | RNAseq |
| APP/PS1 (509) | 558 ♂ | Fibroblast | Newborn; P5 | RNAseq |

For RNAseq of hippocampus samples, four biological replicates were included to ensure robustness and reproducibility. For mass spectrometry of hippocampus samples, two biological replicates were used. Additionally, five biological replicates were employed for RNAseq of fibroblast samples. However, one fibroblast sample (ID 550), subjected to β-secretase inhibition, failed quality control assessment for RNA Integrity Number scoring and thus was not subjected to sequencing. The number of replicates was limited by the availability inherent to a large animal model study. RNA from fibroblasts and hippocampal tissues were sequenced as two independent batches. Fibroblasts were cultured and treated in parallel with pharmacological γ−secretase and β-secretase inhibitors or DMSO.

Experimenters performing mass spectrometry were blinded to group allocation. All other experiments were performed in a nonblinded manner because the researchers were limited. Outliers were identified based on predefined criteria related to gene expression levels. Specifically, two fibroblast samples (ID 606 and 607) were excluded from the analysis due to significantly lower APP expression compared to the other samples. These outliers were identified during initial data processing and were excluded from subsequent analyses to ensure data integrity and consistency in the interpretation of results.

Göttingen minipigs were housed at Research Center Foulum, Department of Animal Science, Aarhus University. All experiments involving the animals are reviewed and approved by the Danish Animal Experiment Inspectorate under the license 2017-15-0201-01251.

### Statistical analysis

Statistical significance between groups was determined using the parametric Student’s t-test within GraphPad Prism software. Data are expressed as mean ± SEM. The overlap of DEGs_Padj_ between groups was assessed using the Fisher’s Exact Test for Count Data in R Studio. Differential gene expression analysis was conducted on RNAseq count tables with the DESeq2 package as indicated in section “RNA Sequencing and Bioinformatics” (Supplementary Material). *P* values were adjusted for multiple testing with the Benjamini-Hochberg procedure which controls false discovery rate (FDR). Each experiment was conducted with more than three biological replicates, and *n* is indicated in the figure legends. Significance is indicated as \**P* < 0.05.

## Supporting information

Suppl all

## List of Supplementary Materials

Materials and Methods

Fig. S1 to Fig S5

Tables S1 to S11

## Acknowledgements

We thank Ida Elisabeth Holm for the dissection of porcine brain samples and for helpful discussions.

## Data and materials availability

All data associated with this study are present in Supplementary Materials or are available at Gene Expression Omnibus (GEO) under records GSE254548 and GSE254571.

