## Supplementary material for "APP and its intracellular domain modulate Alzheimer’s disease risk gene networks in transgenic *APPsw* and *PSEN1M146I* porcine models": Suppl all

**Supplementary materials and methods.**

**Cell cultures.**

*Fibroblasts* – Porcine fibroblasts were obtained from ear biopsies from WT, PS1 and APP/PS1 Göttingen minipigs and cultured in porcine fibroblast medium (Dulbecco's Modified Eagle's Medium (DMEM; Sigma Aldrich), 10% FBS (Life Technologies), 1% P/S (P/S, 10,000 units/mL and 10,000 μg/mL; Life Technologies), 1% Gln (2.92 g/100 mL) and 6.667 ng/mL Fibroblast Growth Factor-basic (bFGF; Life Technologies)) in 5% $CO_2$ in a humidified chamber at 37 $^oC$. Established fibroblasts were split in a 1:3 ratio every 3 days using 0.05% trypsin EDTA (Life Technologies). All fibroblasts used in this study were tested negative for mycoplasma (MYCOPLASMACHECK, Eurofins Genomics).

*Pharmacological treatments* – Porcine fibroblasts were dissociated using 0.05% trypsin EDTA (Life Technologies) and seeded onto 0.1% gelatin-coated culture ware (Fisher Scientific) at a density of $0.33x10^5$ cells/$cm^2$. Fibroblasts were left to settle for 24 hrs in 5% $CO_2$ in a humidified chamber at 37 $^oC$. Fibroblasts were treated with 50 μM LY294002 (in DMSO; Cell Signaling) or 2 μM L-685-458 (γ–secretase inhibitor in DMSO; Merck Millipore), 15 μM β-Secretase Inhibitor IV (in DMSO; Merck Millipore), or DMSO (Sigma Aldrich) for 24 hrs.

**FE65 lentiviral construct, production and transduction.**

*Lentiviral construct* – Lentivirus was used to generate fibroblast lines stably expressing FE65. pLenti-C-Myc-DDK-P2A-Puro carrying open reading frame clone of human FE65 (*APBB1;* NM_001164) was bought from OriGene (#RC202003L3).

*Lentivirus production* – Lentiviral particles were produced using 2nd generation lentiviral packaging vectors. In brief, HEK293T/17 cultured in HEK293T/17 medium were transfected with the plasmid of interest together with packaging plasmid PAX2 and envelope plasmid PMDG2 (psPAX2 and pMD2.G, Addgene plasmid #12260 and #12259) using Lipofectamine™ 3000 Transfection Reagent (Life Technologies). Six hrs post-transfection, the medium was changed to 5% KSR. The supernatant containing the virus particles was collected 24 and 48 hrs post-transfection, filtered through a 0.45 μm filter (Frisenette), concentrated by ultracentrifugation 22500 g for 1 hr and 30 mins at 4 °C and resuspended in PBS, snap-frozen and stored at -80 °C.

*Lentivirus titration and transduction* – Lentiviral titration was performed by quantitative PCR using 7500 Fast real-time PCR system (Applied Biosystems) and primers against the lentiviral backbone (LV2 forward 5’-ACTTGAAAGCGAAAGGAAAC-3’ and reverse 5’-CACCCATCTCTCTCCTTCTAGCC-3’; Sigma Aldrich) and *ALB* (Albumin) (*ALB* forward 5’-TTTGCAGATGTCAGTGAAAGAGA-3’ and reverse 5’-TGGGGAGGCTATAGAAAATAAGG-3’) as an internal control for normalization. Titer was calculated by comparing integrated viral DNA content against a vector reference, pWPXL (pWPXL was a gift from Didier Trono) with known titer determined by Green Fluorescent Protein (GFP) expression in transduced cells. Fibroblasts were infected with lentiviruses at a multiplicity of infection (MOI) 10. The medium was completely changed 16 hrs post-infection.

**Western Blotting.**

*Protein extraction* – Samples were lysed in RIPA lysis buffer (prepared in house; 50 mM Tris HCl, 150 mM NaCl 150 mM, Sodium deoxycholate 0.5%, 0.1% SDS, 1% NP40) containing protease (c0mplete Mini, EDTA-free protease inhibitor cocktail; Roche) and phosphatase (Phosphatase

Inhibitor Cocktail 2 aqueous solution P5726; Sigma Aldrich) inhibitors followed by sonication (Bioruptor®) and centrifuged at 5,000 g for 10 mins at 4 °C. The supernatant was collected, and protein concentration was measured using Bradford protein assay (Bio-Rad) following the manufacturer's protocol.

*Co-immunoprecipitation* – We used anti-c-myc (clone 9E10) tagged magnetic beads (Pierce™ c-Myc Tag Magnetic IP/Co-IP Kit; Thermo Fisher Scientific) to pull down c-Myc tagged proteins in fibroblast lysates following manufacturer's protocol.

*Western blotting* – Proteins were separated by SDS-PAGE on 16.5% Tris/Tricine gels (Bio-Rad) or 10% TGX gels (Bio-Rad) and transferred to nitrocellulose (0.2 µm; Bio-Rad) or PVDF (Bio-Rad) membranes. The membranes were boiled in PBS (Life Technologies) for 5 mins to unmask the epitopes and blocked in 5% skim milk (Difco™) or BSA (Sigma Aldrich), 50 mL TBS-T containing 100 mL 10xTris Buffered Saline (Fisher BioReagents), 900 mL water, 2 mL 100% Tween 20 (Sigma Aldrich)) for 1 hr prior to probing with primary antibody O/N: mouse anti-APP-C-terminus (C1/6.1; 1:2000; Biolegend), rabbit anti-Akt (1:1000, Cell Signaling #9272), rabbit anti-Phospho-Akt (Ser473) (D9E, XP®, 1:2000, Cell Signaling #4060), and mouse anti-beta actin (1:10,000; Abcam; ab6276). Membranes were washed 4x30 mins and applied with horse-radish peroxidase-conjugated secondary antibodies (1:2000; Dako) for 1.5 hrs followed by additional 4x30 mins of washing. Blots were developed using Clarity™ Enhanced Chemiluminescence (Bio-rad) by digital imaging. For reprobing, membranes were stripped using Re-Blot Strong Solution (Calbiochem).

**Immunocytochemistry.**

Fibroblasts plated on coverslips (VWR) were washed twice with PBS (Life Technologies) and fixated in 4% paraformaldehyde (PFA; Santa Cruz Biotechnologies Inc) for 15 mins at 4 $^{o}$C, then washed with PBS (Life Technologies) and distilled water (Life Technologies) and left to dry for 10 mins. Fibroblasts were permeabilized for 10 mins at RT in PBS (Life Technologies) containing 0.25% Triton-X (Sigma Aldrich) (PBT) and blocked for 1 hr at RT in PBT supplemented with 5% donkey serum (Almeco). The primary antibodies were diluted in blocking solution and cells and antibodies were incubated O/N at 4$^{o}$C: rabbit anti-APP-C-terminus (Y188; 1:1000; Abcam) and mouse anti-c-myc (9E10; 1:1000; Thermo Fisher Scientific). Fibroblasts were washed with PBT and blocked for 10 mins. Secondary antibodies conjugated to fluorophores -568 or -488 (Jackson Immunoresearch) were diluted in blocking buffer and applied for 1 hr at RT, then washed and counterstained with 49,6-diamidino-2-phenylindole (DAPI; 1 μg/mL; Sigma) diluted in PBS (Life Technologies). Cells were washed three times with PBS (Life Technologies) and mounted using PVA-DABCO (Sigma Aldrich).

**RNA Sequencing and Bioinformatics.**

*Homogenization and RNA extraction* – Hippocampal tissue from four WT pigs, four PS1 pigs and four APP/PS1 pigs were quickly removed after sacrificing the animals, dissected on ice and immediately submerged into liquid nitrogen and stored at -135 $^{o}$C. Tissues were homogenized in 1 mL cold homogenization buffer (20 μL Thioglycerol per mL of Homogenization solution) using a blender. Homogenates were left to settle on ice. Homogenates (200 μL) were mixed with equal volume Lysis buffer and added to cartridges (Maxwell® 16 LEV simplyRNA Tissue Kit; Promega). DNAse I solution was added before samples were processed on Maxwell ® 16 Instrument. Eluted RNA was stored in -70 $^{o}$C.

Fibroblasts were lysed in TRI Reagent® (Sigma Aldrich), and the mixture was added to equal volumes of ethanol (95-100%). The solution was transferred into Zymo-Spin™ IICR Columns and processed according to the manufacturer's protocol (Direct-zol™ RNA kit; Zymo Research). In-column DNAse I treatment was included. Eluted RNA was stored at -70 °C. All samples preserved RNA integrity as assessed by RIN values scoring 8.0 or higher.

*RNA Sequencing* – Library construction, sequencing and initial data filtering including adaptor removal were performed by BGI Europe Genome Center. Total RNA was subjected to oligo dT based mRNA enrichment. 50 bp single-end read sequencing was performed on BGISEQ-500 (fibroblast samples), and 100 bp paired-end read sequencing was performed on DNBseq platform (hippocampal tissues). More than 20 million clean reads were obtained per sample. Reads were aligned to the porcine genome build Sscrofa11.1 (Ensemble release 92) using HISAT2 aligner (v2.1.0)*(63)*. Transcript quantification was performed using htseq-count (v0.9.1)*(64)* and the read counts were normalized for effective gene length, and sequencing depth to yield Transcripts Per Kilobase Million (TPM). Differentially expressed genes were determined from count tables using DEseq2 (v2.11.40.6)*(65)*. For fibroblast samples, we excluded raw counts not mapping to neuron-expressed genes. Neuron-expressed genes were selected from publicly available data (Brain RNAseq Database; genes were considered as expressed in neurons if count values were 1≤). Heatmaps were constructed by log2 transforming and mean scaling TPM values using the heatmap.2 function in R stats package. For Principal Component Analysis (PCA), PCs were calculated by singular value decomposition of the centered and scaled data matrix using the prcomp function in R stats package.

*Gene set enrichment and gene set association analyses* - Functional analyses of DEGs were performed using Ingenuity Pathway Analysis (IPA). Profiles of transcription factor binding sites (TFBSs) were retrieved from the JASPER database *(66)*. CiiiDER *(67)* was used to identify TFBSs within the promotor sequences (1500 bp upstream and 500 bp downstream of TSS (Homo sapiens GRCh38.94)) in lists of DEGs and background lists consisting of 2500 unregulated genes identified in the respective RNAseq studies. CiiiDER then assessed the enrichment of TFBSs in DEGs compared to the background gene set input using default mapping stringency settings (deficit = 0.15), coverage $p < 0.05$ and site count $p < 0.05$. Gene set association analysis was performed with Multi-marker Analysis of GenoMic Annotation (MAGMA) *(68)* using default settings, based on summary statistics from GWASs on AD *(33)*, Parkinson's *(38)*, Tourette's syndrome *(39)* and psychiatric cross disorder *(40)*.

*URLs* – Galaxy: https://usegalaxy.org/; STRING: https://string-db.org/; CiiiDER: http://ciiider.com/; Brain RNAseq Database: https://www.brainrnaseq.org/; JASPAR: http://jaspar.genereg.net/; RNA expression in brain cell types: http://celltypes.org/brain/; CSEA: http://genetics.wustl.edu/jdlab/csea-tool-2/; Venn plot tool: http://bioinformatics.psb.ugent.be/webtools/Venn/; IPA: https://www.qiagenbioinformatics.com/products/ingenuitypathway-analysis; Subcell Barcode: https://www.subcellbarcode.org/

**LC-MS/MS.**

*Tissue Homogenization* – Hippocampal tissue from two WT pigs, two PS1 pigs and four PS1/APP pigs were prepared as previously described *(29, 69)*. Tissues were homogenized in 1 mL cold lysis buffer (40 mM Tris-HCl, 150 mM KCl, 1% Igepal CA630 detergent, pH 7.4) supplemented with complete protease inhibitor cocktail (Roche), 2 mM EDTA and 1 mM sodium orthovanadate, using a blender. The homogenates were incubated at 4 °C for 1 hr, under rotation, and then centrifuged at 16,000× *g* for 20 min to remove cell debris. The protein concentration of the supernatant was estimated using the 2D Quant Kit (GE Healthcare, Little Chalfont , UK).

*Biotinylated Peptides* – The three synthetic peptides used for peptide pull-down experiments were the same as previously used *(29, 69)*. The peptides were synthesized with a N-terminal biotin moiety followed by a Ttsd linker coupled to a 31 amino acid residue-long synthetic peptide mimicking the last 31 amino acid C-terminal amino acid residues in porcine APP with (p$_{682}$YENPTY$_{687}$) or without phosphorylation at Y$_{682}$ ($_{682}$YENPTY$_{687}$) (APP residue numbering). A peptide with a scrambled sequence was included as a negative control. Peptides were dissolved in dimethyl sulfoxide (DMSO) (10 mg/mL) and stored until further use at −20 °C.

*Peptide Pull-Down* – Biotinylated peptides (5nM) were incubated with 1mg of prewashed Dynabeads M280 Streptavidin (Thermo Fisher Scientific, Waltham, MA, USA) for 3 hrs at 4 °C under rotation. After four washings in 40 mM Tris-HCl, 0.1% BSA, pH 7.4, followed by equilibration of the beads in 40 mM Tris-HCl, 150 mM KCl, 0.1% Igepal CA630 detergent, pH 7.4, dynabeads bound peptides were incubated with 1mg hippocampal lysate for 16 hrs 4 °C. Beads were then washed four times in 40 mM Tris-HCl, 150 mM KCl, 0.1% Igepal CA630, pH 7.4 and two times in 40 mM Tris-HCl, 150 mM KCl, pH 7.4. Bound protein was eluted using 0.1 M glycine, pH 2.8.

*MS Sample Preparation* – Low pH elutes from peptide pull-down experiments were lyophilized and suspended in 20 µL 8 M urea, 0.2 M Tris-HCl, pH 8.3 added 10 mM dithiothreitol and incubated for 30 min. Samples were then incubated with 30 mM iodoacetamide for 30 min at room temperature. Reduced and alkylated samples were then diluted five times before incubated with 0.5 µg trypsin (sequence grade, Sigma-Aldrich Co, St. Louis, MO, USA) overnight at 37 °C. Digested samples were acidified using formic acid and desalted by micro-purification using POROS 50 R2 RP column material (Applied Biosystems, Forster City, CA, USA) packed in gel loader tips. Micro-purified samples were suspended in 0.1% formic acid and kept at −20 °C until LC-MS/MS analysis.

*LC-MS/MS Analysis* – Information dependent acquisition (IDA) analyses were performed on an EASY-nLC II system (Thermo Fisher Scientific, Waltham, MA, USA) connected to a TripleTOF 6600 mass spectrometer (AB SCIEX, Framingham, MA, USA) equipped with a NanoSpray III source (AB SCIEX, Framingham, MA, USA) and operated under Analyst TF 1.6.0 control. Samples were injected and trapped on a C18 pre-column (5 µm, 2 cm × 100 µm I.D., packed in-house). Subsequently, the peptides were eluted to and separated on a 15 cm analytical column (75 µm I.D., packed in-house with RP ReproSil-Pur C18-AQ 3 µm resin (Dr. Maisch GmbH, Ammerbuch-Entringen, Germany) connected in-line to the mass spectrometer. Peptides were eluted at a flow rate of 250 nL/min using a 20 min gradient from 5% to 35% phase B (0.1% formic acid and 100% acetonitrile). Data were acquired using an ion spray voltage of 2.6 kV, a curtain gas of 35, and an interface heater temperature of 150 °C. For IDA experiments, survey scans were acquired in 250 msec and in a mass range of 400-1800 m/z. As many as 25 product ion scans of 25 msec and in the mass range of 100-1800 m/z were collected if exceeding a threshold of 100 counts per second and with a +2 to +5 charge-state. A sweep collision energy setting of 35 ± 15 eV was applied to all precursor ions for collision-induced dissociation. The cycle time of the DIA method was 0.925.

*LC-PRM-MS* – A targeted MS method monitoring Fe65 was established to measure relative quantities of the protein in the peptide pull-down experiment. The PRM method included four peptides (Fe65: DLLLQLEDETLK (+2); NIATSLHEIC[+57]SK (+3); C[+57]LVNGLSLDHSK (+3); and FLSFLAVGR (+2)). The data were acquired using the same instrumental setup as described for the IDA analyses, however, containing the following adjustments of the standard method: the survey scan was reduced to 200 msec, mass range of 400-900 m/z, an inclusion list was added and the MS/MS triggered cps threshold was increased to 2000000 cps. The number of precursor ions to be selected for MS/MS in each cycle was adjusted to number of m/z values in the inclusion list and the mass range of the survey scan was set to ± 50 Da of the highest and lowest precursor ion in the inclusion list. The cycle time was 1.05 sec allowing a minimum of 10 scans across the chromatographic peak required for post-acquisition extracted ion chromatogram.

*Data Processing* – The IDA data were searched against the Swiss-Prot (1,421 sequences) and Tremble (26,516 sequences) Sus scrofa databases (2017_10) using Mascot 2.5 (Matrix Science, London, UK). Trypsin was employed as enzyme allowing one missed cleavage. Carbamidomethyl was entered as a fixed modification, and oxidation of methionine was entered as a variable modification. The mass tolerances of the precursor and product ions were 10 ppm and 0.2 Da, respectively, and the instrument setting was specified as ESI-QUAD-TOF. The significance threshold (p) was set at 0.01 and the expected cut-off at 0.005. The Mascot search results were parsed using MS Data Miner v. 1.3 *(70)*.

The PRM data were imported to Skyline *(71)* and label-free relative quantification was performed at the MS2 level relying on the five most intense transitions. The total intensity for Fe65 was obtained by summing the intensities of transitions monitored for each protein.

**Fig. S1 to S5**

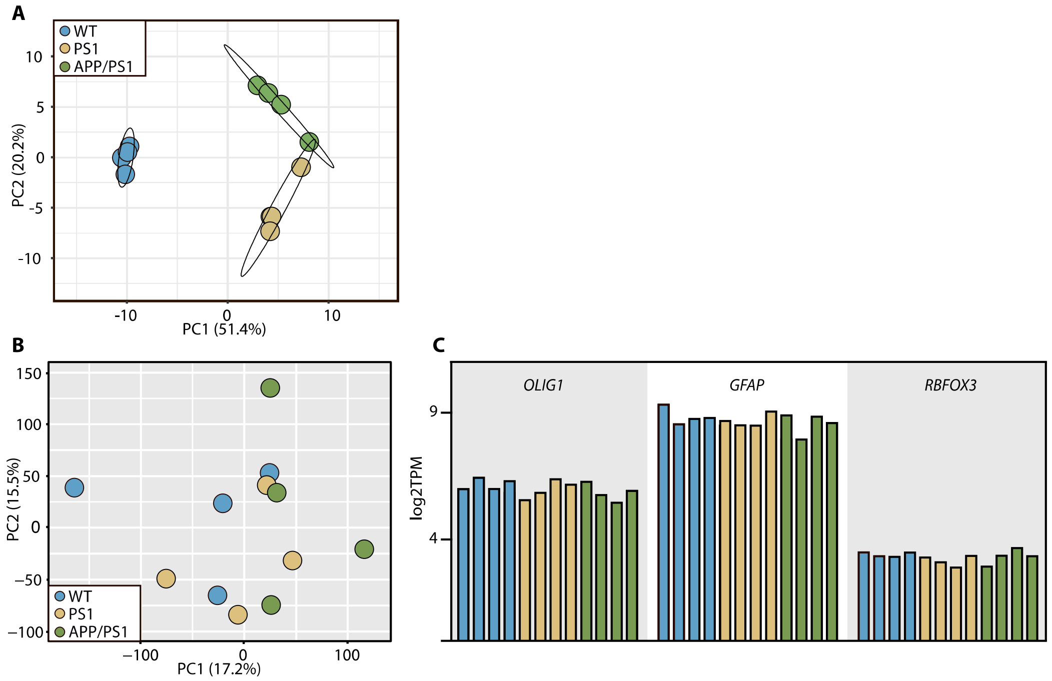


**Fig. S1**: Principal Component Analysis (PCA) plot in (A) using the combined list of DEG$_{Padj}$ identified by comparing WT to PS1 or APP/PS1 hippocampal samples and in (B) using the whole transcriptome of the same samples. (C) Quantification of log2 transformed transcript per million (TPM) values of porcine glia (*OLIG1* and *GFAP*) and neuron (*RBFOX3*) markers. Colors indicate genotype as in A and B.

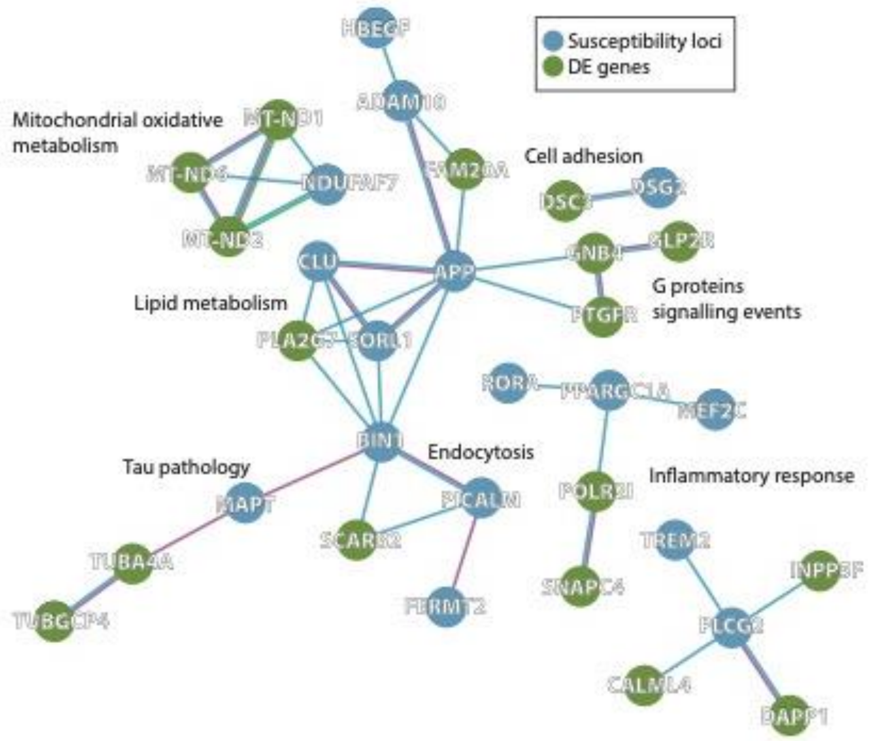


**Fig. S2**: STRING protein-protein interaction network of $DEG_{Padj}$ identified by comparing WT to PS1 or APP/PS1 hippocampal samples (in green) and risk loci reaching genome-wide significance for association with AD (Sims et al., 2020) (in blue). Interaction sources: experimental and database interactions.

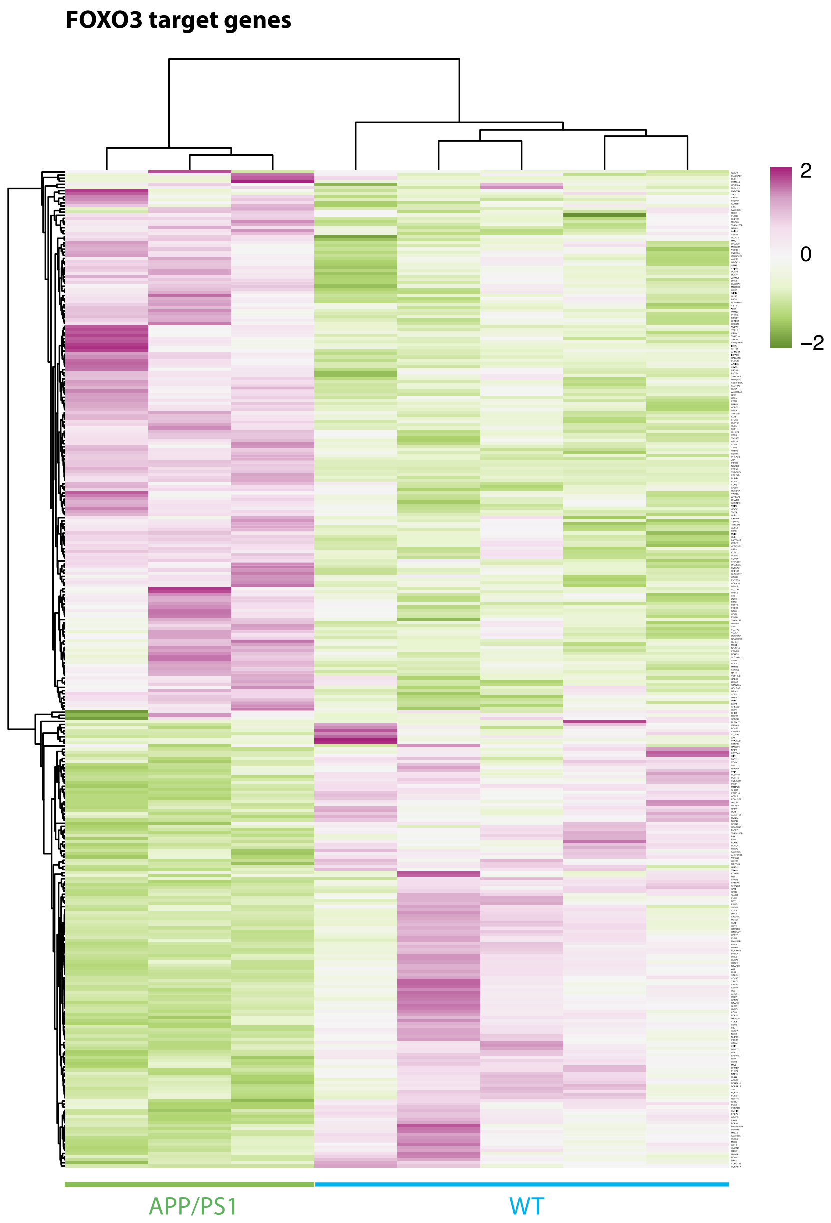


**Fig. S3**: Heatmap of log2 transformed and mean scaled TPM expression values of FOXO3 target genes in APP/PS1 and WT fibroblasts.

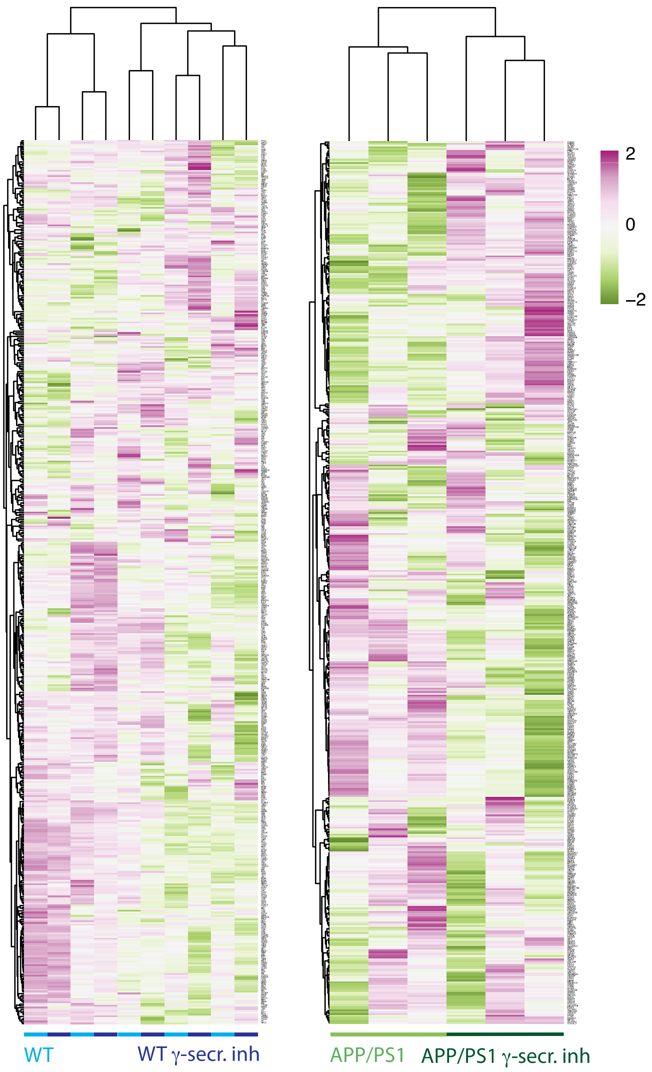


**Fig. S4**: Heatmap of log2 transformed and mean scaled TPM expression values of these genes in WT treated with or without γ– secretase inhibitor and APP/PS1 treated with or without γ–secretase inhibitor.

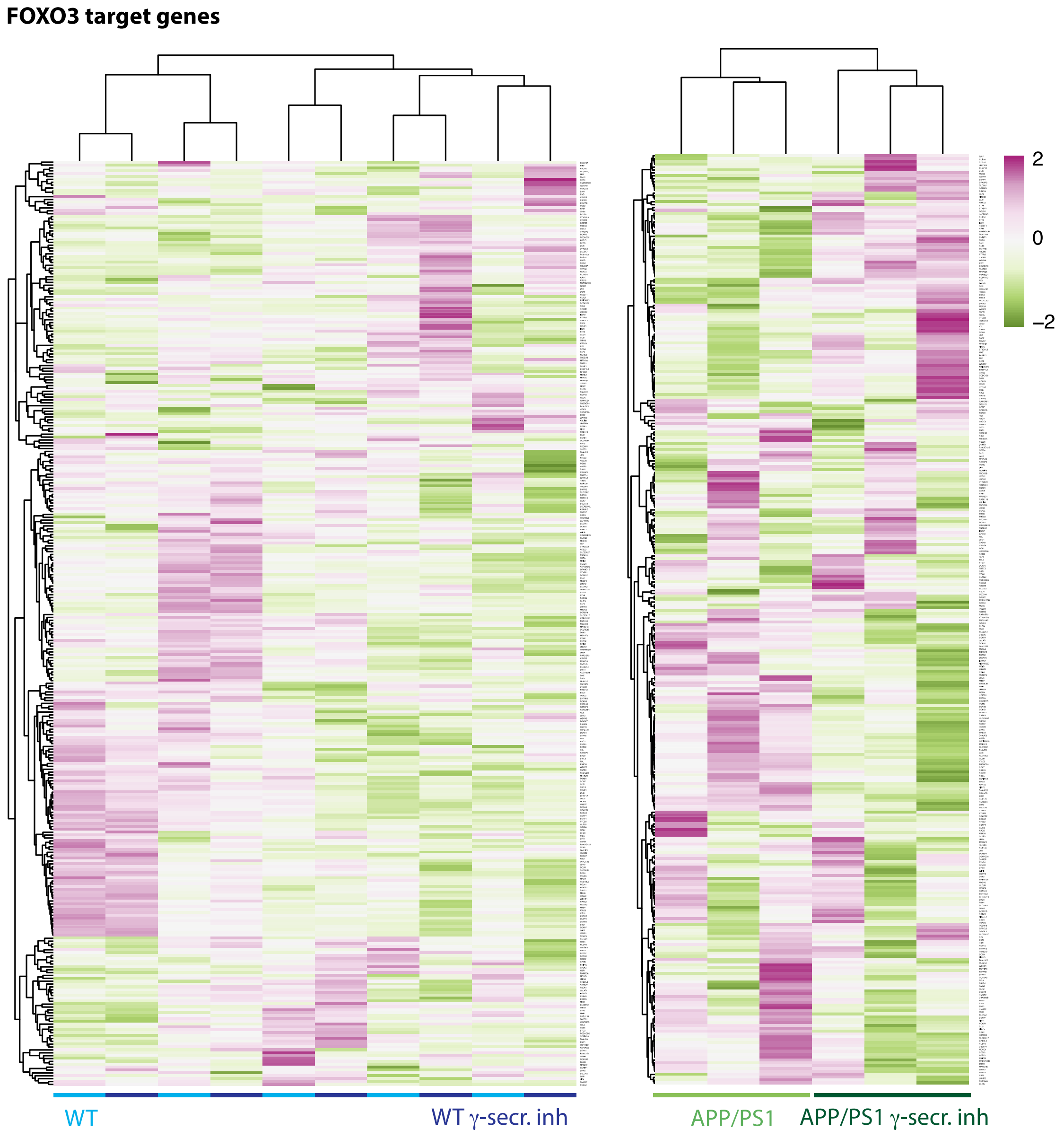


**Fig. S5**: Heatmap of log2 transformed and mean scaled TPM expression values of FOXO3 target genes in fibroblasts from WT treated with or without γ– secretase inhibitor and APP/PS1 treated with or without γ– secretase inhibitor.

**Table S1** | DEG analyses of WT vs PS1 hippocampus (only DEGs$_{Padj}$ with annotated gene symbols are shown). DEGs$_{Padj}$ identified in WT vs APP/PS1 are in red.

| Geneid | Basemean | log2(FC) | Padj |
|---|---|---|---|
| *ABCB9* | 826.10 | -0.44 | 0.0461 |
| *ANGPTL2* | 433.49 | -0.89 | 0.0001 |
| *ARHGAP15* | 83.88 | 0.57 | 0.0389 |
| *ARL4D* | 127.23 | -0.72 | 0.0232 |
| *ATXN7L2* | 490.17 | -0.49 | 0.0025 |
| *BCL2A1* | 37.05 | 0.77 | 0.0004 |
| *BPGM* | 1308.45 | 0.38 | 0.0007 |
| *CCDC34* | 250.02 | 0.54 | 0.0213 |
| *CES3* | 190.83 | -0.9 | 0.0005 |
| *CHMP4C* | 23.07 | -0.69 | 0.0368 |
| *CYP4V2* | 1112.45 | 0.43 | 0.0452 |
| *DAPP1* | 147.17 | 0.81 | 0.0006 |
| *DEAF1* | 3523.78 | 0.66 | 0.0145 |
| *DSC3* | 374.97 | 0.57 | 0.0097 |
| *ETV4* | 211.50 | -0.73 | 0.0001 |
| *FAM163B* | 1310.30 | -0.9 | 0.0000 |
| *FBN2* | 56.02 | -0.81 | 0.0033 |
| *FER* | 908.27 | 0.32 | 0.0034 |
| *FRZB* | 440.44 | 0.5 | 0.0162 |
| *GNB4* | 907.80 | 0.41 | 0.0001 |
| *GRAMD2B* | 2870.79 | 0.31 | 0.0389 |
| *GSG1L* | 235.22 | -0.56 | 0.0166 |
| *KRT23* | 23.97 | 0.96 | 0.0001 |
| *LMOD1* | 178.21 | -0.75 | 0.0010 |
| *LRP3* | 1470.65 | -0.26 | 0.0311 |
| *MMP19* | 111.17 | 0.78 | 0.0032 |
| *NEURL1* | 6555.72 | 0.4 | 0.0213 |
| *PMP2* | 102.95 | 1.56 | 0.0000 |
| *POLR2I* | 537.85 | -0.43 | 0.0244 |
| *PPFIA4* | 4288.01 | -0.46 | 0.0074 |
| *PRDM1* | 60.54 | 0.61 | 0.0376 |
| *PTGFR* | 441.67 | 0.61 | 0.0388 |
| *RNF39* | 76.48 | -0.97 | 0.0001 |
| *SNAPC4* | 1370.82 | -0.3 | 0.0232 |
| *SNCB* | 4611.39 | -0.44 | 0.0389 |
| *SOX18* | 135.29 | -0.64 | 0.0162 |
| *TBL1XR1* | 1063.94 | 0.36 | 0.0431 |
| *TBX6* | 202.66 | -0.47 | 0.0122 |
| *THRSP* | 53.88 | 1.03 | 0.0000 |
| *TIPARP* | 1174.84 | 0.36 | 0.0249 |
| *TMEM145* | 1462.07 | -0.42 | 0.0259 |
| *TMEM14A* | 840.96 | 0.41 | 0.0181 |
| *TOP2A* | 49.39 | 0.63 | 0.0461 |
| *TP53BP1* | 3093.97 | 0.55 | 0.0000 |
| *TUBGCP4* | 2751.40 | 1.47 | 0.0000 |
| *UGT2A3* | 372.81 | 0.85 | 0.0000 |
| *UNC119* | 472.36 | -0.37 | 0.0264 |

**Table S2** | DEG analyses of WT vs APP/PS1 hippocampus. (only DEGs$_{Padj}$ with annotated gene symbols are shown). DEGs$_{Padj}$ identified in WT vs PS1 are in red.

| Geneid | Basemean | log2(FC) | Padj |
|---|---|---|---|
| *ANGPTL2* | 414.49 | -0.76 | 0.0008 |
| *ASAP3* | 711.22 | -0.8 | 0.0000 |
| *ATP8* | 42623.56 | 0.83 | 0.0002 |
| *C19orf54* | 385.62 | 0.6 | 0.0182 |
| *CES3* | 181.34 | -0.74 | 0.0045 |
| *COX2* | 577382.07 | 0.59 | 0.0004 |
| *DEAF1* | 3182.35 | 0.58 | 0.0429 |
| *ECH1* | 436.22 | 0.37 | 0.0498 |
| *FAM20A* | 63.15 | 0.64 | 0.0211 |

| | | | |
|---|---|---|---|
| *GLP2R* | 282.30 | 0.69 | 0.0045 |
| *HIST1H1D* | 263.19 | 0.73 | 0.0051 |
| *IAH1* | 573.27 | -0.39 | 0.0317 |
| *INPP5F* | 38373.94 | 0.89 | 0.0000 |
| *KRT23* | 20.05 | 0.66 | 0.0211 |
| *MNS1* | 445.36 | 0.57 | 0.0419 |
| *MRPL45* | 613.60 | 0.41 | 0.0464 |
| *MT1A* | 64.49 | 0.66 | 0.0240 |
| *ND1* | 185302.07 | 0.66 | 0.0145 |
| *ND2* | 367666.90 | 0.68 | 0.0012 |
| *ND5* | 471868.49 | 0.57 | 0.0006 |
| *ND6* | 137029.07 | 0.76 | 0.0000 |
| *NME5* | 201.78 | -0.64 | 0.0355 |
| *OTOP1* | 73.80 | 0.66 | 0.0159 |
| *P4HA2* | 180.85 | -0.49 | 0.0288 |
| *PLA2G7* | 429.79 | -0.67 | 0.0120 |
| *PMFBP1* | 67.27 | 0.64 | 0.0138 |
| *PMP2* | 78.74 | 0.95 | 0.0000 |
| *SCARB2* | 12121.51 | 0.32 | 0.0475 |
| *TIMD4* | 23.06 | 0.96 | 0.0000 |
| *TNNT1* | 59.74 | 0.63 | 0.0382 |
| *TP53BP1* | 2796.53 | 0.46 | 0.0278 |
| *TTC7A* | 749.48 | 0.6 | 0.0042 |
| *TUBGCP4* | 2652.95 | 1.49 | 0.0000 |
| *UGT2A3* | 363.55 | 0.89 | 0.0000 |
| *VRK3* | 1013.38 | 0.55 | 0.0147 |
| *ZNF503* | 214.00 | -0.5 | 0.0215 |

**Table S3** | IPA pathway analysis of $DEG_{pnom}$ WT vs PS1 hippocampus

| **Ingenuity Canonical Pathways** | **-log(p-value)** | **Ratio** | **z-score** |
|---|---|---|---|
| Calcium Signaling | 3.40 | 5.34E-02 | 0.447 |
| Hepatic Fibrosis / Hepatic Stellate Cell Activation | 3.16 | 5.38E-02 | NaN |
| Mechanisms of Viral Exit from Host Cells | 2.43 | 9.76E-02 | NaN |
| Cellular Effects of Sildenafil (Viagra) | 2.35 | 5.34E-02 | NaN |
| Agrin Interactions at Neuromuscular Junction | 2.33 | 7.14E-02 | -1.342 |
| Tec Kinase Signaling | 2.24 | 4.62E-02 | -0.447 |
| RhoGDI Signaling | 2.02 | 4.23E-02 | 2.449 |
| Calcium Transport I | 2.00 | 2.00E-01 | NaN |
| Agranulocyte Adhesion and Diapedesis | 1.98 | 4.17E-02 | NaN |
| GP6 Signaling Pathway | 1.97 | 5.04E-02 | -1.633 |
| Death Receptor Signaling | 1.84 | 5.43E-02 | -0.447 |
| Fcγ Receptor-mediated Phagocytosis in Macrophages and Monocytes | 1.80 | 5.32E-02 | -1.342 |
| Signaling by Rho Family GTPases | 1.75 | 3.56E-02 | -2.646 |
| UVA-Induced MAPK Signaling | 1.73 | 5.10E-02 | 1 |
| PI3K Signaling in B Lymphocytes | 1.68 | 4.35E-02 | 2.449 |
| Semaphorin Neuronal Repulsive Signaling Pathway | 1.66 | 4.32E-02 | -1.633 |
| Histidine Degradation VI | 1.66 | 1.33E-01 | NaN |
| Axonal Guidance Signaling | 1.63 | 2.83E-02 | NaN |
| FAK Signaling | 1.63 | 4.81E-02 | NaN |
| IGF-1 Signaling | 1.63 | 4.81E-02 | NaN |
| CREB Signaling in Neurons | 1.60 | 2.68E-02 | 1.387 |
| B Cell Receptor Signaling | 1.58 | 3.76E-02 | 1.89 |
| Angiopoietin Signaling | 1.53 | 5.33E-02 | 0 |
| Sulfate Activation for Sulfonation | 1.51 | 5.00E-01 | NaN |
| Leukocyte Extravasation Signaling | 1.50 | 3.63E-02 | -0.378 |
| Epithelial Adherens Junction Signaling | 1.50 | 3.95E-02 | NaN |
| Gap Junction Signaling | 1.45 | 3.54E-02 | NaN |
| Sirtuin Signaling Pathway | 1.41 | 3.09E-02 | 0.378 |
| G Beta Gamma Signaling | 1.38 | 4.10E-02 | 2 |
| L-carnitine Biosynthesis | 1.34 | 3.33E-01 | NaN |
| Inosine-5'-phosphate Biosynthesis II | 1.34 | 3.33E-01 | NaN |
| CXCR4 Signaling | 1.33 | 3.59E-02 | -1 |
| Cardiac Hypertrophy Signaling (Enhanced) | 1.32 | 2.62E-02 | 1.732 |
| Apoptosis Signaling | 1.15 | 4.00E-02 | -2 |
| Regulation of Actin-based Motility by Rho | 1.11 | 3.88E-02 | -2 |
| RhoA Signaling | 0.90 | 3.25E-02 | -2 |

| | | | |
|---|---|---|---|
| 14-3-3-mediated Signaling | 0.87 | 3.15E-02 | 2 |
| ILK Signaling | 0.47 | 2.11E-02 | -2 |

**Table S4** | IPA pathway analysis of $DEG_{pnom}$ WT vs APP/PS1 hippocampus

| Ingenuity Canonical Pathways | -log(p-value) | Ratio | z-score |
|---|---|---|---|
| Mitochondrial Dysfunction | 5.55 | 5.85E-02 | NaN |
| Oxidative Phosphorylation | 5.27 | 7.34E-02 | 2.121 |
| Sirtuin Signaling Pathway | 4.26 | 3.78E-02 | -1.414 |
| SPINK1 General Cancer Pathway | 1.64 | 4.35E-02 | NaN |
| Atherosclerosis Signaling | 1.59 | 3.15E-02 | NaN |
| Oxidized GTP and dGTP Detoxification | 1.59 | 3.33E-01 | NaN |
| Myo-inositol Biosynthesis | 1.37 | 2.00E-01 | NaN |
| Phagosome Maturation | 1.36 | 2.65E-02 | NaN |

**Table S5** | emPAI values for proteins identified in the peptide pull-down experiment (only proteins detected in ≥ one sample is shown)

| Genotype | WT | | | | | | PS1 | | | | | | APP/PS1 (592) | | | | | | APP/PS1 (509) | | | | | |
|---|---|---|---|---|---|---|---|---|---|---|---|---|---|---|---|---|---|---|---|---|---|---|---|---|
| Replicate | 1 | | | 2 | | | 1 | | | 2 | | | 1 | | | 2 | | | 1 | | | 2 | | |
| Peptide | - | | Y | - | | Y | - | | Y | - | | Y | - | | Y | - | | Y | - | | Y | - | | Y |
| Acetyl-CoA carboxylase α | 0.07 | 0.04 | | | 0.24 | | 0.93 | 0.45 | | 1.26 | 0.64 | | 0.64 | 0.09 | | 4.90 | 1.47 | | 0.67 | 0.05 | | 4.79 | 0.50 | 1.43 |
| Serum albumin | 0.59 | 0.39 | 0.81 | 0.39 | 0.39 | 0.30 | 0.69 | 0.69 | 0.81 | 0.81 | 0.69 | 0.69 | 0.81 | 0.39 | 0.14 | 2.07 | | | 0.49 | 0.69 | 0.69 | 0.81 | 0.30 | 0.81 |
| AP complex subunit | 3.89 | 11.67 | 0.89 | 0.89 | 5.71 | 0.89 | 3.89 | 8.22 | 0.89 | 3.89 | 8.22 | 0.89 | 5.72 | 2.56 | 0.89 | 3.89 | 5.72 | | 8.22 | 5.71 | 0.89 | 5.72 | 5.71 | 0.89 |
| AP-2 complex subunit alpha-2 isoform 1 | 4.36 | 5.14 | 0.80 | 0.80 | 3.67 | 1.26 | 3.27 | 4.86 | 1.07 | 4.60 | 5.72 | 1.71 | 4.12 | 2.56 | 0.37 | 3.08 | 4.60 | 0.57 | 4.12 | 6.70 | 1.48 | 3.67 | 4.86 | 1.07 |
| AP complex subunit | | 7.98 | | | 4.74 | | | 7.58 | | 5.86 | 9.27 | | 4.24 | 2.35 | | | 5.56 | | | 6.18 | | | 5.27 | |
| Amyloid beta precursor protein binding family B member 1 | 1.49 | 0.84 | | 0.28 | 0.44 | | 1.20 | 0.44 | | 2.37 | 0.63 | | 1.64 | 0.63 | | 2.17 | 1.20 | 0.20 | 1.20 | 0.63 | 0.13 | 1.99 | 1.07 | 0.20 |
| Amyloid beta precursor protein binding family B member 2 | 1.46 | 0.57 | | | 0.18 | | 1.20 | 0.66 | | 1.46 | 0.48 | | 0.96 | 0.25 | | 1.08 | 1.32 | | 0.66 | 0.32 | | 0.96 | 1.08 | |
| Amyloid protein variant 2 | 0.32 | 0.25 | | 0.18 | 0.25 | | 0.25 | 0.25 | 0.25 | 0.25 | 0.32 | | 0.25 | 0.18 | | 0.25 | 0.25 | | 0.32 | 0.25 | | 0.25 | 0.25 | 0.18 |
| ATP synthase subunit alpha. mitochondrial | 2.54 | 2.27 | 0.17 | 0.88 | 1.38 | 0.37 | 2.54 | 1.20 | 0.61 | 3.49 | 1.79 | | 1.79 | 0.88 | 0.27 | 2.27 | 1.04 | 2.02 | 2.83 | 2.83 | 0.37 | 1.20 | 1.04 | 0.17 |
| ATP synthase subunit beta | 3.06 | 1.23 | | 2.01 | 1.23 | 0.49 | 3.49 | 1.46 | 0.65 | 2.01 | 2.32 | | 3.06 | 0.49 | 0.35 | 3.06 | 2.68 | 5.70 | 3.06 | 2.68 | 0.49 | 1.72 | 2.67 | 1.23 |
| ATP synthase subunit O. mitochondrial | 3.95 | 3.05 | 1.23 | | 3.05 | 2.32 | 3.05 | 5.05 | 2.32 | 3.05 | 3.05 | | 1.72 | 0.82 | | 2.32 | 1.23 | 2.32 | 2.32 | 3.05 | 1.23 | 0.82 | 1.72 | 0.49 |
| ATPase. H+ transportin | 0.51 | 0.73 | | | 0.23 | | 0.32 | 2.00 | | 0.62 | 1.44 | | 0.15 | 0.32 | | | 4.57 | 0.51 | | 2.22 | | 2.44 | 11.71 | 2.69 |

| | | | | | | | | | | | | | | | | | | | | | | | | |
|---|---|---|---|---|---|---|---|---|---|---|---|---|---|---|---|---|---|---|---|---|---|---|---|---|
| **g. lysosomal 70kDa.V1 A** | | | | | | | | | | | | | | | | | | | | | | | | |
| **Branched chain ketoacid dehydrogenase kinase** | | 0.83 | | | 0.66 | | 0.50 | 0.66 | | 0.50 | 0.66 | | 0.50 | 0.35 | | 0.50 | 2.36 | | 0.22 | 0.66 | | 0.83 | 1.48 | |
| **Cbl proto-oncogene. E3 ubiquitin protein ligase** | | 1.10 | | | 0.59 | | | 1.53 | 0.10 | | 1.65 | | | 0.45 | | | 1.65 | | | 0.91 | | | 1.41 | |
| **Cbl proto-oncogene. E3 ubiquitin protein ligase B** | | 0.29 | | | 0.35 | | | 0.35 | | | 0.24 | | | 0.24 | | | 0.67 | | | 0.29 | | | 0.47 | |
| **Cofilin-1** | 2.47 | 2.47 | 1.11 | 2.47 | 1.11 | 1.71 | 1.71 | 2.47 | 1.71 | 2.47 | 3.45 | 1.71 | 1.71 | 0.65 | 1.11 | 1.71 | 1.71 | 8.40 | 2.47 | 6.33 | 1.11 | 3.45 | 3.45 | 4.71 |
| **Clathrin light chain** | 3.42 | 1.69 | 1.69 | 1.69 | 2.45 | 1.69 | 3.41 | 1.69 | 2.45 | 3.41 | 2.45 | 2.45 | 3.41 | 1.10 | 1.69 | 3.41 | 2.45 | 1.10 | 1.69 | 1.69 | 2.45 | 2.45 | 2.45 | 2.45 |
| **Clathrin heavy chain** | 6.64 | 5.43 | 0.19 | 3.14 | 6.27 | 1.94 | 9.50 | 9.00 | 1.19 | 14.92 | 12.75 | 2.09 | 8.76 | 1.66 | 0.55 | 14.16 | 8.07 | 1.09 | 10.58 | 5.43 | 0.76 | 14.54 | 4.16 | 2.75 |
| **2'.3'-cyclic-nucleotide 3'-phosphodiesterase** | 13.47 | 14.97 | 7.82 | 0.22 | 10.87 | 2.62 | 16.63 | 9.75 | 6.24 | 10.87 | 14.97 | 4.38 | 12.10 | 3.00 | 0.64 | 7.82 | 3.87 | 5.56 | 0.64 | 1.00 | 0.22 | 0.22 | | 0.64 |
| **V-crk avian sarcoma virus CT10 oncogene-like** | | 2.04 | | | 2.04 | | | 3.01 | | | 3.01 | | | 0.52 | | | 5.08 | | | 4.29 | | | 3.61 | |
| **Collapsin response mediator protein 1** | 0.50 | 0.50 | | | 0.38 | | 0.76 | 0.62 | | 0.76 | 0.62 | | 0.91 | 0.38 | 0.18 | 1.07 | 0.50 | 1.63 | 1.07 | 0.91 | | 0.50 | 0.91 | 0.38 |
| **Dynamin 1** | 0.97 | 10.24 | 0.34 | 0.16 | 10.80 | 0.88 | 3.70 | 20.08 | 1.51 | 3.27 | 23.38 | 2.04 | 2.88 | 4.18 | 0.16 | 0.40 | 9.20 | 0.47 | 0.10 | 13.32 | 1.07 | | 1.90 | |
| **Dihydropyrimidinase like 2** | 2.82 | 3.34 | 0.38 | 0.14 | 1.96 | 0.78 | 2.15 | 2.58 | 1.44 | 3.34 | 3.62 | 0.67 | 3.07 | 0.67 | 0.21 | 2.15 | 2.15 | 5.36 | 2.58 | 3.93 | 0.21 | 1.02 | 2.15 | 1.15 |
| **Elongation factor 1-alpha** | 0.45 | 0.32 | 0.75 | 2.69 | 0.32 | | 0.21 | 0.32 | | 0.32 | 0.59 | 0.45 | 0.59 | 0.21 | | 0.32 | 0.32 | 0.75 | 0.59 | 0.45 | 0.21 | 0.45 | | |
| **Epidermal growth factor receptor pathway substrate 15 like 1** | | 0.42 | | | 0.53 | | | 1.69 | | 0.53 | 3.74 | | 0.42 | 1.03 | | | 0.89 | | 0.15 | 0.33 | | 1.34 | 1.18 | |
| **Tyrosine-protein kinase Fyn** | | 2.18 | | | 1.94 | | | 3.01 | | | 6.43 | 0.36 | | 1.16 | | | 3.33 | | | 4.05 | | | 2.44 | |
| **Glyceraldehyde-3-phosphate dehydrogenase** | 10.91 | 9.45 | 8.18 | 2.23 | 12.57 | 4.45 | 9.45 | 7.05 | 4.45 | 14.46 | 12.57 | 9.45 | 10.91 | | | 7.05 | 5.20 | 10.91 | 9.45 | 7.05 | 3.78 | 3.78 | 2.68 | 2.23 |
| **Growth factor receptor bound protein 2** | | 32.76 | 0.45 | | 22.31 | 0.45 | | 39.64 | 2.04 | | 57.86 | 2.04 | | 8.23 | | | 39.64 | 2.66 | | 32.77 | 2.04 | | 22.31 | 0.74 |

| | | | | | | | | | | | | | | | | | | | | | | | | |
|---|---|---|---|---|---|---|---|---|---|---|---|---|---|---|---|---|---|---|---|---|---|---|---|---|
| **GULP. engulfment adaptor PTB domain containing 1** | 5.85 | 8.21 | | | 6.94 | | 8.21 | 6.94 | 0.34 | 13.35 | 15.64 | 0.34 | 8.21 | 5.85 | | 11.38 | 6.94 | | 5.85 | 5.85 | | 5.85 | 2.79 | |
| **Histone H1.2-like protein** | 1.40 | 0.55 | 0.93 | 0.93 | 0.55 | | 0.93 | 0.93 | 0.55 | 1.40 | 0.55 | 0.55 | 0.93 | 1.40 | 0.55 | 1.40 | | 1.40 | 0.93 | | 0.55 | 0.93 | 0.55 | |
| **Hemoglobin subunit alpha** | 3.67 | 3.67 | 1.52 | 2.43 | 2.43 | 1.52 | 3.67 | 1.52 | 3.67 | 3.67 | 0.85 | 2.43 | 3.67 | 1.52 | | 5.36 | 2.43 | 15.02 | 7.65 | 5.36 | 2.43 | 2.43 | 2.43 | 1.52 |
| **17beta-estradiol dehydrogenase** | 1.88 | 1.27 | 0.12 | | 0.12 | | 1.41 | 0.60 | 0.19 | 1.14 | 1.56 | 0.19 | 0.80 | 0.34 | | 0.91 | 0.42 | | 1.41 | 1.02 | 0.51 | 1.27 | 0.70 | 0.42 |
| **Inositol polyphosphate-5-phosphatase D** | | 0.18 | | | 0.28 | | | 0.28 | | | 0.51 | | | 0.09 | | | 0.93 | | | 0.13 | | | 0.45 | |
| **Potassium voltage-gated channel subfamily A regulatory beta subunit 2** | 2.97 | 6.44 | 0.29 | | 1.73 | | 2.09 | 3.50 | 0.29 | 0.87 | 4.11 | | 0.46 | 1.12 | | 2.50 | 4.11 | | 0.29 | 2.50 | | | 0.65 | |
| **Histone H2B** | 1.73 | 4.33 | 1.73 | 9.42 | 1.73 | 0.95 | 0.95 | | | 2.82 | 1.73 | 2.82 | 4.33 | | | 4.33 | 4.33 | 4.33 | 1.73 | | 2.82 | 2.82 | 1.73 | 1.73 |
| **Myelin basic protein** | 8.40 | 11.05 | 6.33 | 6.33 | 24.44 | 8.40 | 8.40 | 14.46 | 6.33 | 8.40 | 8.40 | 8.40 | 14.46 | 8.40 | 2.47 | 8.40 | 8.40 | 11.05 | 6.33 | 6.33 | 8.40 | 6.33 | 6.33 | 8.40 |
| **NUMB like. endocytic adaptor protein** | 2.66 | 5.06 | | | 4.25 | | 3.23 | 5.51 | 0.33 | 2.93 | 6.00 | | 2.40 | 1.21 | | 2.93 | 4.64 | | 1.21 | 4.25 | | 2.40 | 5.51 | 0.43 |
| **Papilin. proteoglycan like sulfated glycoprotein** | 1.36 | 2.74 | | | 2.28 | | 1.36 | 2.99 | 0.22 | 1.36 | 2.99 | | 1.21 | 0.69 | | 1.36 | 3.27 | | 0.59 | 2.50 | 0.22 | 1.36 | 3.27 | |
| **Propionyl-CoA carboxylase alpha subunit** | 5.62 | 3.83 | 2.53 | 0.88 | 2.53 | 2.53 | 8.07 | 1.57 | 2.53 | 3.83 | 2.53 | 2.53 | 5.62 | 0.88 | | 2.53 | 0.88 | | 5.62 | 3.83 | 5.62 | 0.88 | | |
| **Propionyl-CoA carboxylase beta chain. mitochondrial** | 8.33 | 5.78 | 3.93 | 0.75 | 2.58 | 3.55 | 6.95 | 2.88 | 5.26 | 6.34 | 3.20 | 5.78 | 5.26 | 1.05 | 0.27 | 2.58 | 0.89 | 0.75 | 9.10 | 6.95 | 9.10 | 0.61 | | |
| **Phosphatidylinositol 4.5-bisphosphate 3-kinase catalytic subunit β isoform 2** | | 0.77 | | | 0.64 | | | 1.06 | | | 1.80 | | | 0.58 | | | 1.22 | | | 0.91 | | | 0.16 | |
| **Phosphoinositide-3-kinase. regulatory subunit 1 (Alpha)** | | 6.17 | | | 2.86 | | | 3.84 | 0.12 | | 4.73 | 0.18 | | 1.08 | | | 4.41 | | | 5.41 | 0.18 | | 4.73 | |
| **Peptidyl-prolyl cis-trans isomerase A** | 1.81 | 0.68 | 0.68 | 9.19 | 1.17 | | 0.68 | 0.68 | | 1.81 | 1.17 | | 1.17 | 0.68 | 1.17 | 1.81 | 1.17 | 12.19 | 1.81 | 12.19 | | 1.17 | 2.63 | 1.17 |

| | | | | | | | | | | | | | | | | | | | | | | | | |
|---|---|---|---|---|---|---|---|---|---|---|---|---|---|---|---|---|---|---|---|---|---|---|---|---|
| **Pyrroline-5-carboxylate reductase** | 20.23 | 17.47 | 8.23 | 4.29 | 15.07 | 7.03 | 31.2 | 17.47 | 11.18 | 35.98 | 20.22 | 17.47 | 27.01 | 7.03 | 0.74 | 31.18 | 23.38 | 11.18 | 20.22 | 20.22 | 9.60 | 12.99 | 9.60 | 9.60 |
| **RALBP1 associated Eps domain containing 1** | 0.72 | 0.72 | | | 0.39 | | 0.46 | 0.63 | | 1.26 | 1.66 | 0.18 | 0.82 | 0.82 | | 0.39 | 0.92 | | 0.54 | 0.54 | | 0.72 | 0.92 | |
| **Son of sevenless-like 1** | | 0.10 | | | 0.36 | | | 0.59 | | | 1.03 | | | 0.40 | | | 0.64 | | | 0.54 | | | 0.13 | |
| **Tubulin beta chain** | 5.625 | 4.815 | 2.39 | 3.085 | 9.005 | 2.52 | 9.415 | 7.98 | 6.165 | 11.55 | 19.205 | 3.265 | 9.865 | 2.71 | | 14.745 | 12.065 | 19.04 | 7.635 | 4.94 | 2.775 | 14.97 | 6.7 | 8.895 |
| **Uncharacterized protein** | 3.08 | 2.37 | 0.29 | 0.29 | 0.78 | 0.56 | 3.63 | 1.29 | 1.45 | 1.96 | 1.29 | 0.47 | 1.45 | 0.47 | 0.14 | 2.59 | 1.78 | 0.67 | 2.82 | 1.96 | 1.29 | 0.78 | 1.61 | 0.38 |
| **Uncharacterized protein** | 3.55 | 4.54 | 0.93 | 3.86 | 3.26 | | 3.86 | 4.92 | 0.81 | 4.19 | 4.54 | 1.51 | 3.55 | 1.35 | | 3.55 | 5.75 | 3.86 | 2.49 | 4.19 | 0.59 | 4.54 | 4.54 | 2.99 |
| **Uncharacterized protein** | 0.95 | 0.95 | 0.95 | 7.71 | 1.71 | | 1.30 | 2.79 | 0.65 | 3.47 | 2.21 | | 2.21 | 1.30 | 0.65 | 2.21 | 1.71 | 5.24 | 2.79 | 2.21 | | 1.30 | 2.21 | |
| **Uncharacterized protein** | 4.25 | 2.98 | 2.02 | 2.98 | 5.92 | 1.29 | 4.25 | 2.98 | 2.02 | 2.98 | 5.92 | 1.29 | 2.98 | 1.29 | | 4.25 | 4.25 | 5.92 | 4.25 | 4.25 | 1.29 | 8.12 | 14.85 | 14.85 |
| **Uncharacterized protein** | 0.95 | 0.77 | | 1.59 | 7.12 | 1.36 | 2.45 | 4.05 | 0.46 | 5.71 | 8.83 | 0.77 | 7.93 | 2.13 | 0.61 | 3.59 | 5.10 | 7.12 | 0.61 | 0.77 | | 0.33 | 0.21 | |
| **Uncharacterized protein** | 3.46 | 1.71 | 0.65 | 1.71 | 2.31 | | 5.64 | 2.65 | 1.99 | 6.34 | 3.04 | 3.04 | 1.45 | 0.49 | 0.22 | 3.46 | 3.46 | 1.22 | 3.93 | 3.04 | 1.22 | 3.93 | 3.04 | 2.65 |
| **Uncharacterized protein** | 11.78 | 23.74 | | | 15.96 | | 14.43 | 21.51 | 8.63 | 14.44 | 23.74 | 9.58 | 15.96 | 4.47 | | | 21.51 | | 11.78 | 23.74 | 5.01 | 8.63 | 14.43 | |
| **Uncharacterized protein** | 0.31 | 0.10 | | | 0.20 | | 0.51 | 0.31 | | 1.60 | 1.37 | | 0.81 | 0.58 | | 0.44 | 0.10 | | 0.44 | 0.20 | | 0.81 | 0.26 | |
| **Uncharacterized protein** | | 1.84 | | | 1.38 | | | 0.89 | | | 1.68 | | | 0.89 | | | 4.36 | | | 4.36 | | | 2.37 | |

**Table S6** | DEG analysis WT vs APP/PS1 Fibroblasts. (only DEGs$_{Padj}$ p <0.001 with annotated gene symbols are shown)

| Gene Symbol | Base Mean | log2FC | Padj |
|---|---:|---:|---:|
| PTPRS | 17486.7839562719 | 3.65338818968378 | 0 |
| TUBGCP4 | 1565.62913945407 | 2.16831479848235 | 2.20772510402354E-91 |
| PRPF4B | 5424.72637833283 | 1.96387566505023 | 5.31632976700695E-67 |
| APP | 25695.520351932 | 2.10079023499336 | 4.60074223206535E-40 |
| WSCD2 | 2505.56194097954 | 2.99546626938026 | 1.12841429625838E-30 |
| PSEN1 | 2019.21350958658 | 1.19736840029285 | 4.17671056481975E-29 |
| OXTR | 362.922635814836 | 1.42501109175495 | 6.69717335851161E-15 |
| SH2B3 | 1471.38924126412 | -0.982672811408915 | 6.58475122147317E-13 |
| NFIL3 | 415.68986752538 | 0.927010088836242 | 2.48559270484054E-11 |
| CDCA7 | 597.925989035448 | -1.09243413475053 | 4.96933806565073E-10 |

| | | | |
|---|---|---|---|
| GAS7 | 1079.86949120952 | -1.45241370192759 | 8.92268438480977E-10 |
| SRGN | 67.4100301327539 | 1.66434501181294 | 1.14982801392596E-08 |
| PRICKLE1 | 527.572049884281 | -0.869118896525669 | 1.14982801392596E-08 |
| FDFT1 | 2363.16780096816 | 0.666626865134606 | 1.99386405778038E-08 |
| MTUS2 | 98.3150073253097 | 1.55646714503695 | 2.76410899908115E-08 |
| AXL | 5181.54533437047 | -0.699117439833537 | 2.76410899908115E-08 |
| FAM20C | 3549.0803056945 | 1.25034349144013 | 3.40308245905199E-08 |
| KLF6 | 427.775061797663 | 0.728978145317779 | 3.58276832938564E-08 |
| TCP11L2 | 974.465921750027 | 1.01165277501765 | 5.20815600265009E-08 |
| RASL11B | 197.21365326157 | 1.37836506073996 | 1.21444808804728E-07 |
| EIF2AK3 | 1180.33562443643 | 0.76563054257181 | 2.0042128140808E-07 |
| OSTM1 | 1476.89521303419 | 0.628946192246534 | 2.23753311958268E-07 |
| SYT11 | 704.668326244813 | 1.09053164966218 | 2.23753311958268E-07 |
| SLC1A4 | 1433.65015635609 | 1.04377543911079 | 2.23753311958268E-07 |
| MEST | 84.4199047342013 | 1.43802510347586 | 2.46899469498848E-07 |
| IMPAD1 | 7143.50590566626 | 0.834135680148473 | 4.39639471567373E-07 |
| PDIA3 | 18167.1049521697 | 0.423101400853097 | 1.03192104102292E-06 |
| NUDT6 | 326.655876042473 | 0.868606277815982 | 1.03192104102292E-06 |
| EHBP1L1 | 2066.7145206188 | -0.661440307647499 | 2.20344756910525E-06 |
| FAM169A | 26.4096220801454 | 1.42239681856105 | 2.20344756910525E-06 |
| PDE4D | 1421.46385030883 | 1.06341957592725 | 2.20344756910525E-06 |
| MEF2A | 4740.46307445374 | 0.633335137875903 | 2.20344756910525E-06 |
| B3GNT7 | 37.2960383083411 | 1.37948972784702 | 4.27121220635023E-06 |
| NNAT | 585.701140595003 | 1.02549738337771 | 6.74408731792766E-06 |
| HTRA3 | 1407.7502676913 | -1.1496170672375 | 7.48384633915749E-06 |
| PCOLCE2 | 1647.19281811525 | -0.796569779822902 | 7.48384633915749E-06 |
| CUL1 | 2518.35799760679 | 0.576822404470331 | 7.48384633915749E-06 |
| MAZ | 1827.84955744287 | -0.602785670241191 | 8.37851617919944E-06 |
| PTP4A3 | 71.8058243488526 | -1.02944182408838 | 8.37851617919944E-06 |
| LRIG3 | 653.666227327809 | 0.98558543854119 | 8.41878252204441E-06 |
| LFNG | 72.3607706080139 | -1.2211415540368 | 0.00001350625765789 |
| NT5C2 | 4521.66615154151 | 0.628408366746502 | 0.0000176357282093535 |
| BMPR2 | 3834.50543549975 | 0.572622314102767 | 0.00001993543695227 |
| CADM1 | 77.9700826167334 | -1.12833425545159 | 0.0000370057327281632 |
| CENPA | 546.179879147644 | -0.693340375271315 | 0.0000414513936471575 |
| NOTCH1 | 584.736757237582 | -1.25161733313503 | 0.0000435758267936889 |
| DLL4 | 57.5448477552632 | 1.25560559867342 | 0.0000456079031953129 |
| GALK2 | 6820.7466459472 | 0.769088063337812 | 0.000055595171935721 |
| NCAPD2 | 1069.85650597138 | -0.702564576064683 | 0.0000686324056970413 |
| LIMD1 | 725.850213306491 | -0.594784556257948 | 0.0000757171361993433 |

| | | | |
|---|---|---|---|
| HMGN2 | 7378.32843762642 | -0.613177761639617 | 0.0000883364080320503 |
| BCAR3 | 548.46185970534 | -0.921080481629877 | 0.0000883364080320503 |
| SIPA1L2 | 180.807585670788 | 1.06538031982009 | 0.0000883364080320503 |
| NBEA | 444.987570160805 | 1.00246180211175 | 0.0000883364080320503 |
| STXBP1 | 586.832259884001 | 0.836804056058991 | 0.000107627509115185 |
| RDH10 | 624.765410678724 | 0.999538785431184 | 0.000107627509115185 |
| AKT3 | 1220.16177735065 | 0.60726319491939 | 0.000110989266623052 |
| SLC1A1 | 39.4476049878402 | 1.170636148428 | 0.000111740401012738 |
| DLC1 | 2453.49101827877 | 0.469907363838105 | 0.000126686998837153 |
| NEK2 | 320.876420462818 | -0.733328875541786 | 0.000126686998837153 |
| MCM4 | 2057.40922417632 | -0.817904312353766 | 0.000148885239087181 |
| PRKAR1B | 38.112985947569 | 1.19090144167117 | 0.000148885239087181 |
| LRRTM4 | 66.2274276963525 | -1.20111039243001 | 0.000148885239087181 |
| UNG | 473.944325884598 | -0.696834860214338 | 0.000149562382729262 |
| SLC18A2 | 198.019327329526 | 0.692874187244494 | 0.000149562382729262 |
| POU3F2 | 19.8182298228801 | 1.17593952315248 | 0.000155951938417087 |
| PCK2 | 601.292999468168 | -0.868363306847104 | 0.000170632279806339 |
| ACSL5 | 919.900851421099 | -0.711293308326125 | 0.000176598673889904 |
| POLE4 | 628.36172456965 | -0.610129744791284 | 0.000176598673889904 |
| RABEP1 | 1802.67301639132 | 0.549982480331391 | 0.000199621993783705 |
| DIO2 | 56.815020590066 | 1.17538541589283 | 0.000207284234173285 |
| RAP2B | 1497.41429509962 | -0.506937814524609 | 0.00023183100159322 |
| GFPT1 | 1859.02868924202 | 0.425669945265106 | 0.000289308359787306 |
| UBE2C | 1411.45903067228 | -0.697918462062664 | 0.000290333788502404 |
| SFR1 | 107.633816752642 | -0.896550752437497 | 0.000306859251766109 |
| PRKACB | 1037.42233184414 | 0.657447634913685 | 0.000306859251766109 |
| PABPC1 | 26118.26145995 | -0.425527575499419 | 0.00031738229949203 |
| TRMT112 | 800.739440004773 | -0.480922443170616 | 0.00031738229949203 |
| AHCY | 1460.90728618398 | -0.618212832678447 | 0.000330971654883965 |
| CLGN | 515.478015534275 | 1.09743176285685 | 0.000359548488837844 |
| C2CD2 | 993.7449652538 | 0.695162932933195 | 0.00045988294792988 |
| HS6ST1 | 1961.03970337231 | 0.778498314228653 | 0.000470437316359221 |
| DKK3 | 4403.82257431116 | 0.82753932496742 | 0.000524346573259871 |
| RNF145 | 1758.03608429521 | 0.57970458826295 | 0.000524346573259871 |
| TIMELESS | 341.162100192729 | -0.722381160819008 | 0.000528654953530646 |
| TMEM135 | 567.437970906226 | 0.607787866054604 | 0.000528654953530646 |
| LYAR | 528.978324509204 | -0.734993708586601 | 0.000561833624249034 |
| TCF19 | 322.963609952888 | -0.824638222250339 | 0.000575674464786321 |
| NDRG2 | 689.763472821261 | 1.03483147437005 | 0.000589766625073638 |
| LMNB1 | 2080.89631870016 | -0.638265890365655 | 0.000617669741344521 |

| | | | |
|---|---|---|---|
| EEF1A2 | 692.594544871185 | -1.08557000104364 | 0.000650017339586226 |
| DNMT1 | 2948.14285231653 | -0.565208039616489 | 0.000686809371249962 |
| NCAPH | 706.075342575485 | -0.694571914389529 | 0.000725716261308819 |
| HSPE1 | 1442.82246892702 | -0.56679522334564 | 0.000766248200139358 |
| ACSL3 | 908.631120769733 | 0.493253858964602 | 0.000816808320123714 |
| DNAJB9 | 660.749279609599 | 0.681153930880691 | 0.00082612823309612 |
| FBLN1 | 423.753070574813 | 1.07157530206982 | 0.000829580649884789 |
| SLC36A4 | 527.905934698425 | 0.654394723377072 | 0.000851119269689351 |
| PGM2 | 620.306035037923 | 0.466296860352803 | 0.000914809152287899 |
| ZFAND5 | 1771.43051325122 | 0.435479414448282 | 0.000914809152287899 |
| FBXO5 | 343.317030572818 | -0.756883009971209 | 0.000918154115000261 |
| KAT2A | 540.53979800703 | -0.454390971548201 | 0.000918154115000261 |
| PER3 | 448.612910417662 | -0.561802641918139 | 0.000922169951292323 |
| SSRP1 | 4948.276153101 | -0.363111172926384 | 0.000965495891972367 |
| MRTO4 | 726.53538819054 | -0.445518255705041 | 0.000965495891972367 |

**Table S7** | IPA pathway analysis of $DEG_{Padj}$ WT vs APP/PS1 fibroblasts

| **Ingenuity Canonical Pathways** | **-log(p-** | **Ratio** | **z-score** |
|---|---|---|---|
| Cell Cycle Control of Chromosomal Replication | 10.90 | 0.25 | -3.207 |
| NER Pathway | 7.24 | 0.14 | -2.714 |
| Antiproliferative Role of TOB in T Cell Signaling | 4.71 | 0.18 | 1.89 |
| FAT10 Cancer Signaling Pathway | 4.22 | 0.15 | 2.646 |
| Molecular Mechanisms of Cancer | 4.22 | 0.06 | NaN |
| Cell Cycle: G1/S Checkpoint Regulation | 3.97 | 0.12 | 0.82 |
| Pancreatic Adenocarcinoma Signaling | 3.84 | 0.09 | -0.38 |
| Unfolded protein response | 3.67 | 0.13 | 1.633 |
| Leptin Signaling in Obesity | 3.67 | 0.11 | NaN |
| Role of NFAT in Cardiac Hypertrophy | 3.54 | 0.07 | 1.941 |
| Granzyme B Signaling | 3.47 | 0.25 | -1 |
| Mismatch Repair in Eukaryotes | 3.47 | 0.25 | NaN |
| Kinetochore Metaphase Signaling Pathway | 3.42 | 0.09 | -2.449 |
| Senescence Pathway | 3.39 | 0.06 | 1.5 |
| RAN Signaling | 3.36 | 0.24 | -2 |
| Chronic Myeloid Leukemia Signaling | 3.35 | 0.09 | NaN |
| Endoplasmic Reticulum Stress Pathway | 2.99 | 0.19 | 1 |
| p53 Signaling | 2.84 | 0.08 | 0.45 |
| Coronavirus Pathogenesis Pathway | 2.74 | 0.07 | 2.53 |
| Cyclins and Cell Cycle Regulation | 2.69 | 0.09 | -1.342 |
| Semaphorin Neuronal Repulsive Signaling Pathway | 2.63 | 0.07 | -1 |
| Estrogen-mediated S-phase Entry | 2.63 | 0.15 | -1 |
| Aldosterone Signaling in Epithelial Cells | 2.58 | 0.06 | 0 |
| DNA Double-Strand Break Repair by Homologous Recombination | 2.50 | 0.21 | NaN |
| Pyridoxal 5'-phosphate Salvage Pathway | 2.49 | 0.09 | -0.82 |
| Glioblastoma Multiforme Signaling | 2.44 | 0.06 | -0.71 |
| Sperm Motility | 2.40 | 0.05 | 1.667 |
| Cell Cycle: G2/M DNA Damage Checkpoint Regulation | 2.37 | 0.10 | -1 |
| Sphingosine-1-phosphate Signaling | 2.37 | 0.07 | -0.71 |
| CMP-N-acetylneuraminate Biosynthesis I (Eukaryotes) | 2.34 | 0.40 | NaN |
| Amyloid Processing | 2.33 | 0.10 | NaN |
| Salvage Pathways of Pyrimidine Ribonucleotides | 2.23 | 0.07 | -1.134 |
| UVA-Induced MAPK Signaling | 2.23 | 0.07 | -0.82 |
| CREB Signaling in Neurons | 2.21 | 0.05 | 0.33 |
| Inhibition of Angiogenesis by TSP1 | 2.20 | 0.12 | NaN |

| | | | |
|---|---|---|---|
| PTEN Signaling | 2.17 | 0.06 | -0.38 |
| Neuropathic Pain Signaling In Dorsal Horn Neurons | 2.16 | 0.07 | 0.38 |
| P2Y Purigenic Receptor Signaling Pathway | 2.15 | 0.06 | 0.71 |
| Protein Kinase A Signaling | 2.13 | 0.04 | 1.698 |
| GADD45 Signaling | 2.11 | 0.16 | NaN |
| Cell Cycle Regulation by BTG Family Proteins | 2.07 | 0.11 | 0 |
| Notch Signaling | 2.07 | 0.11 | 1 |
| Regulation Of The Epithelial Mesenchymal Transition By Growth Factors Pathway | 2.05 | 0.05 | 0 |
| CDP-diacylglycerol Biosynthesis I | 2.04 | 0.15 | NaN |
| Docosahexaenoic Acid (DHA) Signaling | 2.03 | 0.11 | NaN |
| PPARα/RXRα Activation | 2.02 | 0.05 | -0.38 |
| Adipogenesis pathway | 2.02 | 0.06 | NaN |
| Regulation of the Epithelial-Mesenchymal Transition Pathway | 2.00 | 0.05 | NaN |
| Colorectal Cancer Metastasis Signaling | 1.98 | 0.05 | -0.33 |
| Glioma Signaling | 1.97 | 0.06 | -1 |
| PI3K Signaling in B Lymphocytes | 1.95 | 0.06 | -0.38 |
| Phosphatidylglycerol Biosynthesis II (Non-plastidic) | 1.92 | 0.14 | NaN |
| Adrenomedullin signaling pathway | 1.92 | 0.05 | 0.33 |
| Hereditary Breast Cancer Signaling | 1.91 | 0.06 | NaN |
| Th1 and Th2 Activation Pathway | 1.87 | 0.05 | NaN |
| Apelin Endothelial Signaling Pathway | 1.87 | 0.06 | 0.38 |
| PXR/RXR Activation | 1.85 | 0.08 | NaN |
| Role of Tissue Factor in Cancer | 1.83 | 0.06 | NaN |
| Apelin Pancreas Signaling Pathway | 1.81 | 0.09 | -2 |
| D-myo-inositol (1.4.5)-Trisphosphate Biosynthesis | 1.77 | 0.12 | NaN |
| Role of Oct4 in Mammalian Embryonic Stem Cell Pluripotency | 1.74 | 0.09 | NaN |
| PAK Signaling | 1.70 | 0.06 | -0.82 |
| AMPK Signaling | 1.69 | 0.05 | 1.633 |
| Melatonin Signaling | 1.68 | 0.07 | 1.342 |
| Synaptogenesis Signaling Pathway | 1.68 | 0.04 | 0 |
| Lanosterol Biosynthesis | 1.66 | 1.00 | NaN |
| VEGF Signaling | 1.66 | 0.06 | -1 |
| Non-Small Cell Lung Cancer Signaling | 1.66 | 0.07 | -1 |
| GPCR-Mediated Integration of Enteroendocrine Signaling Exemplified by an L Cell | 1.66 | 0.07 | 0.45 |
| Endocannabinoid Neuronal Synapse Pathway | 1.64 | 0.05 | -0.38 |
| GDP-glucose Biosynthesis | 1.63 | 0.18 | NaN |
| Synaptic Long Term Potentiation | 1.62 | 0.05 | 0.38 |
| Melanoma Signaling | 1.62 | 0.08 | -1 |
| Superpathway of Cholesterol Biosynthesis | 1.60 | 0.10 | NaN |
| Cellular Effects of Sildenafil (Viagra) | 1.59 | 0.05 | NaN |
| IGF-1 Signaling | 1.57 | 0.06 | 0.45 |
| Glucose and Glucose-1-phosphate Degradation | 1.56 | 0.17 | NaN |
| Glycogen Degradation II | 1.56 | 0.17 | NaN |
| BER pathway | 1.56 | 0.17 | NaN |
| IL-7 Signaling Pathway | 1.55 | 0.06 | NaN |
| Human Embryonic Stem Cell Pluripotency | 1.53 | 0.05 | NaN |
| Estrogen Receptor Signaling | 1.52 | 0.04 | 0.28 |
| Th2 Pathway | 1.51 | 0.05 | -1.89 |
| Gap Junction Signaling | 1.51 | 0.05 | NaN |
| Fatty Acid Activation | 1.49 | 0.15 | NaN |
| Cholesterol Biosynthesis I | 1.49 | 0.15 | NaN |
| Cholesterol Biosynthesis II (via 24.25-dihydrolanosterol) | 1.49 | 0.15 | NaN |
| Cholesterol Biosynthesis III (via Desmosterol) | 1.49 | 0.15 | NaN |
| Fatty Acid β-oxidation I | 1.49 | 0.09 | NaN |
| Neuroinflammation Signaling Pathway | 1.48 | 0.04 | 1.732 |
| Ovarian Cancer Signaling | 1.47 | 0.05 | 0 |
| CNTF Signaling | 1.44 | 0.07 | -2 |
| Role of CHK Proteins in Cell Cycle Checkpoint Control | 1.44 | 0.07 | NaN |
| Glycogen Degradation III | 1.44 | 0.14 | NaN |
| Huntington's Disease Signaling | 1.43 | 0.04 | -0.82 |
| GPCR-Mediated Nutrient Sensing in Enteroendocrine Cells | 1.43 | 0.05 | 1.633 |
| Cancer Drug Resistance By Drug Efflux | 1.42 | 0.07 | NaN |

| | | | |
|---|---|---|---|
| G-Protein Coupled Receptor Signaling | 1.41 | 0.04 | NaN |
| Calcium Signaling | 1.41 | 0.04 | 0.38 |
| BMP signaling pathway | 1.41 | 0.06 | 1 |
| Cardiac Hypertrophy Signaling | 1.40 | 0.04 | 1.265 |
| Nucleotide Excision Repair Pathway | 1.39 | 0.09 | NaN |
| Cardiac Hypertrophy Signaling (Enhanced) | 1.38 | 0.03 | 2.668 |
| Endometrial Cancer Signaling | 1.37 | 0.07 | -1 |
| Insulin Secretion Signaling Pathway | 1.37 | 0.04 | 0 |
| Epoxysqualene Biosynthesis | 1.36 | 0.50 | NaN |
| Taurine Biosynthesis | 1.36 | 0.50 | NaN |
| NF-κB Signaling | 1.35 | 0.04 | 0 |
| IL-2 Signaling | 1.35 | 0.07 | -1 |
| p38 MAPK Signaling | 1.34 | 0.05 | 1.342 |
| Regulation of IL-2 Expression in Activated and Anergic T Lymphocytes | 1.34 | 0.06 | NaN |
| Relaxin Signaling | 1.32 | 0.05 | 1 |
| Factors Promoting Cardiogenesis in Vertebrates | 1.32 | 0.05 | 1.89 |
| Th1 Pathway | 1.29 | 0.05 | 0 |
| Phospholipases | 1.28 | 0.06 | 1 |
| γ-linolenate Biosynthesis II (Animals) | 1.28 | 0.12 | NaN |
| Mitochondrial L-carnitine Shuttle Pathway | 1.28 | 0.12 | NaN |
| Inhibition of Matrix Metalloproteases | 1.27 | 0.08 | NaN |
| RhoA Signaling | 1.27 | 0.05 | 0 |
| Netrin Signaling | 1.26 | 0.06 | 2 |
| Melanocyte Development and Pigmentation Signaling | 1.25 | 0.05 | -0.447 |
| Mitotic Roles of Polo-Like Kinase | 1.24 | 0.06 | NaN |
| NRF2-mediated Oxidative Stress Response | 1.24 | 0.04 | 1 |
| Regulation of eIF4 and p70S6K Signaling | 1.23 | 0.04 | NaN |
| EIF2 Signaling | 1.22 | 0.04 | -1.342 |
| Neuregulin Signaling | 1.22 | 0.05 | -0.447 |
| TGF-β Signaling | 1.22 | 0.05 | 1 |
| Mechanisms of Viral Exit from Host Cells | 1.22 | 0.07 | NaN |
| 14-3-3-mediated Signaling | 1.21 | 0.05 | 0 |

**Table S8** | IPA Upstream regulator analysis of $DEG_{Padj}$ WT vs APP/PS1 fibroblasts

| **Upstream Regulator** | **Expr Log Ratio** | **Predicted Activation State** | **Activation z-score** | **p-value of overlap** |
|---|---|---|---|---|
| TP53 | | | 1.749 | 0.00000000000000000185 |
| TBX2 | | Inhibited | -3.769 | 0.000000000000067 |
| E2F1 | 0.525 | Inhibited | -2.517 | 0.0000000000000796 |
| E2F4 | 0.46 | | | 0.0000000000146 |
| MYC | 0.365 | Inhibited | -3.644 | 0.000000000115 |
| CDKN2A | | Activated | 2.933 | 0.00000000475 |
| FOXO3 | -0.512 | | 1.472 | 0.0000000204 |
| MITF | | Inhibited | -2.985 | 0.0000000276 |
| YY1 | 0.056 | | | 0.000000042 |
| RB1 | -0.02 | Activated | 2.708 | 0.0000000695 |
| CREB1 | 0.057 | | 1.017 | 0.000000123 |
| TP73 | | | 1.61 | 0.000000251 |
| FOXM1 | 0.713 | Inhibited | -3.387 | 0.000000635 |
| E2F3 | 0.114 | Inhibited | -2.538 | 0.000000663 |
| CCND1 | 0.014 | Inhibited | -3.203 | 0.000000737 |
| NUPR1 | -0.214 | Activated | 2.785 | 0.00000526 |
| HTT | 0.101 | | 0.447 | 0.00000742 |
| SP1 | 0.126 | | -1.015 | 0.00000771 |
| RBL1 | 0.494 | Activated | 2.608 | 0.0000091 |
| POU5F1 | | | -1.826 | 0.0000104 |
| ZBTB17 | -0.05 | | | 0.000016 |
| MYOD1 | | | -0.359 | 0.0000164 |
| KLF4 | 0.693 | | -1.406 | 0.0000246 |
| TP63 | | | -1.091 | 0.0000392 |
| RRP1B | 0.276 | | | 0.0000406 |
| LEF1 | | | 0.577 | 0.000041 |
| CLOCK | -0.102 | | 0 | 0.0000936 |
| GLI1 | | | -1.562 | 0.0000991 |
| CEBPA | 0.232 | | 1.211 | 0.000112 |

| | | | | |
|---|---|---|---|---|
| TRRAP | 0.124 | | | 0.000158 |
| Tcf7 | | | 0.535 | 0.000165 |
| MAX | 0.097 | | -1.981 | 0.000185 |
| HNF4A | | | -0.44 | 0.000212 |
| E2F2 | | | | 0.000241 |
| CCNE1 | | | -1.982 | 0.000258 |
| TCF3 | 0.085 | | 0.956 | 0.00036 |
| E2F6 | 0.11 | | 1.915 | 0.00049 |
| PAX3 | | | | 0.000579 |
| TOB1 | -0.474 | | 1.633 | 0.000715 |
| MYBL2 | | | -1.387 | 0.000715 |
| ZBTB7A | 0.007 | | -0.6 | 0.000819 |
| BRCA1 | | | -0.931 | 0.000851 |
| WT1 | | | -0.8 | 0.000989 |
| TEAD2 | -0.035 | | | 0.000991 |
| GATA1 | | Activated | 2.853 | 0.00116 |
| MEF2D | -0.003 | Activated | 2.433 | 0.00121 |
| CNBP | 0.146 | | | 0.00123 |
| NFYB | 0 | | | 0.00134 |
| DDIT3 | -0.101 | | -1.586 | 0.00147 |
| NIPBL | -0.03 | | | 0.00151 |
| RUVBL2 | 0.331 | | | 0.00166 |
| SREBF1 | -0.029 | | 1.813 | 0.00198 |
| KLF5 | -0.687 | | -1.767 | 0.00213 |
| XBP1 | -0.264 | Activated | 2.348 | 0.00273 |
| GMNN | 0.753 | | 0.816 | 0.00275 |
| SOX2 | 0.055 | | -1.131 | 0.00292 |
| CTNNB1 | -0.177 | | 1.152 | 0.00306 |
| MED1 | 0.054 | Inhibited | -2.121 | 0.00343 |
| NFYA | 0.11 | | -0.562 | 0.00359 |
| EGR3 | 0.157 | | -0.277 | 0.00372 |
| EGR2 | 0.248 | | 1.803 | 0.00391 |
| SP3 | 0.024 | | -1.441 | 0.00392 |
| SP4 | -0.003 | | | 0.00393 |
| SMAD3 | -0.407 | | 0.554 | 0.00407 |
| TCF4 | 0.139 | | 0 | 0.00495 |
| CEBPB | 0.055 | | 0.321 | 0.00501 |
| MESP1 | | | | 0.00518 |
| BHLHA15 | | | | 0.00518 |
| ONECUT1 | | | | 0.00523 |
| ASCL1 | | | -0.447 | 0.0055 |
| EP300 | -0.05 | | 0.914 | 0.00565 |
| PPARGC1B | 0.548 | Activated | 2.4 | 0.00578 |
| SMARCB1 | 0.092 | Activated | 2.646 | 0.00594 |
| SOX1 | | | 0.816 | 0.00618 |
| CREB3 | -0.205 | | | 0.0073 |
| MYB | | | -1.942 | 0.00779 |
| NFATC2 | -0.075 | | 1.405 | 0.00803 |
| CREBZF | 0 | | | 0.00853 |
| FOXP3 | | Activated | 2.236 | 0.00871 |
| CREM | -0.289 | | 1.134 | 0.00891 |
| SNAI1 | | | -0.564 | 0.00891 |
| KLF11 | -0.254 | | 0.447 | 0.00916 |
| SOX3 | | | 0.816 | 0.00957 |
| NONO | 0.117 | | | 0.00987 |
| ZNF100 | | | | 0.00996 |
| ZNF85 | | | | 0.00996 |
| ZNF254 | | | | 0.00996 |
| ZNF431 | | | | 0.00996 |
| ZNF43 | | | | 0.00996 |
| ZNF429 | | | | 0.00996 |
| ELL2 | -0.442 | Activated | 2.236 | 0.00997 |
| SOX4 | -0.052 | | -0.475 | 0.0104 |
| NOTCH3 | 0.121 | | -0.811 | 0.0115 |
| HSF1 | 0.224 | | -1.044 | 0.0115 |
| YAP1 | -0.245 | | -1.067 | 0.0118 |
| PSIP1 | -0.017 | | | 0.0131 |
| ZNF91 | | | | 0.0131 |
| TAL1 | | | -1.414 | 0.0135 |

| | | | | |
|---|---|---|---|---|
| RUVBL1 | 0.13 | | | 0.0136 |
| TWIST1 | | | -1.744 | 0.014 |
| PRDM5 | 0.272 | | | 0.0141 |
| ATF6 | -0.135 | | 0.611 | 0.0149 |
| BMI1 | | | | 0.0158 |
| FOXC1 | | | | 0.0164 |
| MSGN1 | | | | 0.0166 |
| TFCP2L1 | | | | 0.0166 |
| EPAS1 | -0.295 | | 1.168 | 0.0166 |
| E2F5 | -0.012 | | | 0.0183 |
| FOXO1 | -0.131 | | -1.929 | 0.0183 |
| PCGF2 | 0.079 | | 1.206 | 0.0186 |
| NOTCH1 | 1.252 | | 0.306 | 0.0196 |
| EZH2 | 0.367 | | -1.144 | 0.0199 |
| PYCARD | | | | 0.0204 |
| PSMD10 | 0.569 | | | 0.0204 |
| ARNT | 0.246 | | | 0.0211 |
| ZNF687 | | | | 0.0226 |
| PMF1/PMF1-BGLAP | 0.595 | | | 0.0226 |
| PRDM14 | | | | 0.0226 |
| BARX1 | | | | 0.0226 |
| CAND1 | -0.27 | | | 0.0226 |
| DMRT1 | | | | 0.0229 |
| NFKBIA | -0.381 | | 1.265 | 0.0244 |
| SIX1 | | | | 0.0248 |
| TRPS1 | 0.012 | | 1 | 0.0263 |
| HNF1A | | | 1.067 | 0.0276 |
| ATF4 | 0.048 | | 0.454 | 0.0278 |
| ETV6 | 0.19 | | | 0.029 |
| MYCN | 0.121 | | -1.086 | 0.0311 |
| MDM2 | 0.075 | | -1.949 | 0.0319 |
| STAT4 | | Activated | 2.46 | 0.0328 |
| ATN1 | 0.003 | | | 0.033 |
| KLF6 | -0.729 | | 0.271 | 0.0335 |
| NKX3-1 | | | | 0.0337 |
| JARID2 | 0.02 | | | 0.0338 |
| PA2G4 | | | | 0.0338 |
| SOX10 | -0.161 | | | 0.0352 |
| KDM5B | -0.369 | Activated | 2.635 | 0.0353 |
| CREBBP | 0.098 | | -1.154 | 0.0357 |
| TSHZ3 | 0.008 | | -0.849 | 0.0357 |
| PPARGC1A | 0.132 | | 0.353 | 0.0364 |
| GATA6 | | | 0.134 | 0.0369 |
| GON4L | | | | 0.0389 |
| MRTFB | 0.284 | | 1 | 0.0389 |
| NFE2L2 | -0.142 | Activated | 3.191 | 0.0422 |
| APBB1 | -0.381 | | | 0.0442 |
| LBH | -0.305 | | | 0.0448 |
| NAT8 | | | | 0.0448 |
| ZNF326 | | | | 0.0448 |
| POU6F1 | | | | 0.0448 |
| ECD | 0.214 | | | 0.0448 |
| FOXC2 | | | -1.964 | 0.0468 |
| CBX4 | -0.117 | | | 0.0498 |
| Cux1 | | | | 0.0498 |

**Table S9** | CiiiDER analysis of $DEG_{Padj}$ WT vs APP/PS1 fibroblasts

| **Transcription Factor ID** | **Transcription Factor Name** | **Total No. Search Genes** | **No. Transcription Factor Search Genes** | **Total No. Background Genes** | **No. Transcription Factor Background Genes** | **Gene Representation** | **Gene P-Value (P<0.04)** | **Average Log2 Proportion Bound** | **Log2 Enrichment** |
|---|---|---|---|---|---|---|---|---|---|
| MA1127.1 | FOSB::JUN | 481 | 2 | 8120 | 245 | Down | 1.2E- | -6.3E+00 | -2.5E+00 |
| MA1139.1 | FOSL2::JUN | 481 | 3 | 8120 | 229 | Down | 1.3E- | -6.1E+00 | -2.0E+00 |
| MA0749.1 | ZBED1 | 481 | 3 | 8120 | 3 | Up | 3.1E- | -9.1E+00 | 4.1E+00 |

| | | | | | | | | | |
|---|---|---|---|---|---|---|---|---|---|
| MA1644.1 | NFYC | 481 | 118 | 8120 | 1563 | Up | 5.4E- | -2.2E+00 | 3.5E-01 |
| MA0060.3 | NFYA | 481 | 118 | 8120 | 1563 | Up | 5.4E- | -2.2E+00 | 3.5E-01 |
| MA1136.1 | FOSB::JUNB | 481 | 4 | 8120 | 232 | Down | 5.6E- | -5.9E+00 | -1.6E+00 |
| MA1511.1 | KLF10 | 481 | 112 | 8120 | 1479 | Up | 6.5E- | -2.3E+00 | 3.6E-01 |
| MA0605.2 | ATF3 | 481 | 9 | 8120 | 350 | Down | 6.6E- | -5.1E+00 | -1.1E+00 |
| MA0030.1 | FOXF2 | 481 | 79 | 8120 | 993 | Up | 8.5E- | -2.8E+00 | 4.3E-01 |
| MA1528.1 | NFIX | 481 | 14 | 8120 | 109 | Up | 9.4E- | -5.6E+00 | 1.2E+00 |
| MA1140.2 | JUNB | 481 | 1 | 8120 | 128 | Down | 1.0E- | -7.2E+00 | -2.3E+00 |
| MA0614.1 | Foxj2 | 481 | 202 | 8120 | 2939 | Up | 1.1E- | -1.4E+00 | 2.2E-01 |
| MA0613.1 | FOXG1 | 481 | 202 | 8120 | 2939 | Up | 1.1E- | -1.4E+00 | 2.2E-01 |
| MA0850.1 | FOXP3 | 481 | 202 | 8120 | 2940 | Up | 1.1E- | -1.4E+00 | 2.2E-01 |
| MA1527.1 | NFIC | 481 | 12 | 8120 | 85 | Up | 1.1E- | -5.9E+00 | 1.3E+00 |
| MA0482.2 | GATA4 | 481 | 58 | 8120 | 1326 | Down | 1.3E- | -2.8E+00 | -4.3E-01 |
| MA0036.3 | GATA2 | 481 | 58 | 8120 | 1326 | Down | 1.3E- | -2.8E+00 | -4.3E-01 |
| MA0037.3 | GATA3 | 481 | 408 | 8120 | 6514 | Up | 1.3E- | -2.8E-01 | 8.1E-02 |
| MA0609.2 | CREM | 481 | 7 | 8120 | 284 | Down | 1.3E- | -5.4E+00 | -1.2E+00 |
| MA0593.1 | FOXP2 | 481 | 107 | 8120 | 1436 | Up | 1.4E- | -2.3E+00 | 3.4E-01 |
| MA0047.3 | FOXA2 | 481 | 107 | 8120 | 1436 | Up | 1.4E- | -2.3E+00 | 3.4E-01 |
| MA1683.1 | FOXA3 | 481 | 107 | 8120 | 1436 | Up | 1.4E- | -2.3E+00 | 3.4E-01 |
| MA0852.2 | FOXK1 | 481 | 107 | 8120 | 1436 | Up | 1.4E- | -2.3E+00 | 3.4E-01 |
| MA1103.2 | FOXK2 | 481 | 107 | 8120 | 1436 | Up | 1.4E- | -2.3E+00 | 3.4E-01 |
| MA0481.3 | FOXP1 | 481 | 107 | 8120 | 1436 | Up | 1.4E- | -2.3E+00 | 3.4E-01 |
| MA0031.1 | FOXD1 | 481 | 107 | 8120 | 1440 | Up | 1.4E- | -2.3E+00 | 3.3E-01 |
| MA0042.2 | FOXI1 | 481 | 107 | 8120 | 1440 | Up | 1.4E- | -2.3E+00 | 3.3E-01 |
| MA0849.1 | FOXO6 | 481 | 107 | 8120 | 1440 | Up | 1.4E- | -2.3E+00 | 3.3E-01 |
| MA0848.1 | FOXO4 | 481 | 107 | 8120 | 1440 | Up | 1.4E- | -2.3E+00 | 3.3E-01 |
| MA0157.2 | FOXO3 | 481 | 107 | 8120 | 1440 | Up | 1.4E- | -2.3E+00 | 3.3E-01 |
| MA1129.1 | FOSL1::JUN | 481 | 1 | 8120 | 119 | Down | 1.5E- | -7.2E+00 | -2.2E+00 |
| MA1104.2 | GATA6 | 481 | 58 | 8120 | 1319 | Down | 1.5E- | -2.8E+00 | -4.2E-01 |
| MA1489.1 | FOXN3 | 481 | 41 | 8120 | 470 | Up | 1.7E- | -3.8E+00 | 5.7E-01 |
| MA0765.2 | ETV5 | 481 | 95 | 8120 | 1987 | Down | 1.8E- | -2.2E+00 | -3.0E-01 |
| MA0750.2 | ZBTB7A | 481 | 124 | 8120 | 2504 | Down | 1.9E- | -1.8E+00 | -2.5E-01 |
| MA1475.1 | CREB3L4 | 481 | 14 | 8120 | 432 | Down | 1.9E- | -4.6E+00 | -8.2E-01 |
| MA0028.2 | ELK1 | 481 | 38 | 8120 | 921 | Down | 2.1E- | -3.4E+00 | -5.1E-01 |
| MA0763.1 | ETV3 | 481 | 38 | 8120 | 921 | Down | 2.1E- | -3.4E+00 | -5.1E-01 |
| MA0838.1 | CEBPG | 481 | 11 | 8120 | 84 | Up | 2.1E- | -6.0E+00 | 1.2E+00 |
| MA1126.1 | FOS::JUN | 481 | 0 | 8120 | 79 | Down | 2.3E- | -8.3E+00 | -3.2E+00 |
| MA0156.2 | FEV | 481 | 52 | 8120 | 1179 | Down | 2.3E- | -3.0E+00 | -4.1E-01 |
| MA1604.1 | Ebf2 | 481 | 61 | 8120 | 766 | Up | 2.5E- | -3.2E+00 | 4.4E-01 |
| MA0076.2 | ELK4 | 481 | 128 | 8120 | 2555 | Down | 2.6E- | -1.8E+00 | -2.4E-01 |
| MA1657.1 | ZNF652 | 481 | 11 | 8120 | 353 | Down | 2.6E- | -5.0E+00 | -8.7E-01 |
| MA0596.1 | SREBF2 | 481 | 94 | 8120 | 1946 | Down | 2.7E- | -2.2E+00 | -2.9E-01 |
| MA1637.1 | EBF3 | 481 | 80 | 8120 | 1065 | Up | 3.2E- | -2.8E+00 | 3.5E-01 |
| MA0834.1 | ATF7 | 481 | 2 | 8120 | 132 | Down | 3.5E- | -6.8E+00 | -1.7E+00 |
| MA1133.1 | JUN::JUNB | 481 | 4 | 8120 | 180 | Down | 3.5E- | -6.1E+00 | -1.2E+00 |
| MA0656.1 | JDP2 | 481 | 2 | 8120 | 133 | Down | 3.5E- | -6.8E+00 | -1.7E+00 |
| MA0840.1 | Creb5 | 481 | 2 | 8120 | 133 | Down | 3.5E- | -6.8E+00 | -1.7E+00 |
| MA1145.1 | FOSL2::JUN | 481 | 12 | 8120 | 366 | Down | 3.8E- | -4.9E+00 | -8.0E-01 |

**Table S10 |** DEG analysis APP/PS1 vs APP/PS1 γ-secretase inhibited. (only DEGs$_{Padj}$ p <0.01 with annotated gene symbols are shown)

| Gene Symbol | Base Mean | log2FC | Padj |
|---|---|---|---|
| NUPR1 | 4005.0258 | 0.82869196 | 3.64E-12 |
| TSPAN18 | 931.397343 | 0.94176833 | 7.34E-10 |
| CSF1 | 4797.96791 | 0.71945714 | 5.00E-07 |
| VEGFB | 608.934102 | 0.75020763 | 1.21E-05 |
| BDKRB2 | 1176.58723 | 0.56507707 | 1.67E-05 |
| LAMA5 | 2099.33366 | 0.64959264 | 1.67E-05 |
| SIRT2 | 842.461933 | 0.6326368 | 3.03E-05 |
| GCLM | 1658.06796 | -0.6427019 | 4.78E-05 |

| | | | |
|---|---|---|---|
| MEGF6 | 785.526244 | 0.67744819 | 0.00018167 |
| UAP1L1 | 723.93502 | 0.65978007 | 0.00018167 |
| DNM1 | 970.183207 | 0.66165442 | 0.00024686 |
| SLC6A8 | 947.413024 | 0.69173061 | 0.00024686 |
| PLEKHO1 | 715.054737 | 0.55255525 | 0.00024686 |
| KPNA2 | 2378.55526 | -0.480613 | 0.00024686 |
| NBL1 | 5564.14067 | 0.58274967 | 0.00060006 |
| KANK2 | 5667.20435 | 0.48301912 | 0.00073307 |
| IFI30 | 622.531611 | 0.6386147 | 0.00092647 |
| VAT1 | 3033.21754 | 0.52622826 | 0.00101354 |
| LTBP3 | 2595.96518 | 0.59641766 | 0.00101511 |
| HOMER3 | 1231.88001 | 0.49956318 | 0.00102457 |
| TIPARP | 742.097532 | -0.4685925 | 0.00109293 |
| CSPG4 | 2050.06669 | 0.61803818 | 0.00127398 |
| KIF11 | 1420.78554 | -0.5285543 | 0.00128813 |
| GNPDA1 | 519.198855 | 0.55265048 | 0.00128813 |
| MMP14 | 20316.196 | 0.5200976 | 0.00137948 |
| HSPA5 | 15289.6862 | -0.6405357 | 0.00137948 |
| CLU | 15748.3994 | 0.42700397 | 0.00139856 |
| NECAB3 | 697.875301 | 0.60425288 | 0.00139856 |
| COL18A1 | 11868.7998 | 0.48370394 | 0.0014929 |
| PLD3 | 3530.42825 | 0.43410165 | 0.00152769 |
| ISLR | 4056.59171 | 0.52437812 | 0.00165039 |
| ATP1B1 | 3854.95524 | -0.5245375 | 0.00173607 |
| MANF | 1261.09136 | -0.5415996 | 0.00203648 |
| SNX8 | 477.335803 | 0.52342161 | 0.00211002 |
| GAS6 | 811.338186 | 0.56588061 | 0.0023932 |
| CCNB2 | 510.402634 | -0.5050995 | 0.0024537 |
| MYEF2 | 733.810163 | -0.4761783 | 0.00268753 |
| CLCN6 | 565.562184 | 0.53203811 | 0.00272596 |
| TOLLIP | 545.340203 | 0.52475528 | 0.00272596 |
| SQSTM1 | 6588.60098 | 0.51073176 | 0.00362281 |
| ODC1 | 1676.91687 | -0.415375 | 0.00398875 |
| PACS2 | 1656.00986 | 0.47248794 | 0.00468868 |
| DOCK6 | 424.309513 | 0.48331569 | 0.00468868 |
| PDE3B | 1389.58818 | -0.5781257 | 0.0047649 |
| WDR81 | 590.318165 | 0.48481014 | 0.00876798 |
| ARSA | 394.274544 | 0.55361273 | 0.00921893 |

**Table S11 |** IPA Upstream regulator analysis of $DEG_{Padj}$ APP/PS1 vs APP/PS1 γ-secretase inhibited (top 50)

| Upstream Regulator | Expr Log Ratio | Predicted Activation State | Activation z-score | p-value of overlap |
|---|---|---|---|---|
| TP53 | | Activated | 2.730 | 1.32E-12 |
| NFKBIA | 0.068 | | -0.577 | 2.22E-07 |
| TCF4 | -0.02 | | -0.277 | 2.57E-07 |
| TP73 | | | 1.322 | 3.44E-07 |
| XBP1 | -0.29 | Inhibited | -2.835 | 1.17E-06 |
| FOXO1 | 0.369 | | 0.478 | 1.85E-06 |
| FOS | 0.492 | | 0.044 | 3.02E-06 |
| FOXO3 | 0.036 | | 1.002 | 3.27E-06 |
| SMARCA4 | 0.017 | | 1.732 | 3.36E-06 |
| MYC | 0.171 | Inhibited | -2.424 | 4.17E-06 |
| CREB3 | -0.155 | | -1.980 | 3.15E-05 |
| NFYA | -0.144 | | 0.728 | 3.54E-05 |
| NFYC | 0.165 | | | 7.36E-05 |
| SPDEF | | | -0.277 | 8.38E-05 |
| ZFAT | 0.027 | | | 1.08E-04 |
| PPARGC1A | 0.045 | | 1.841 | 1.16E-04 |
| TCF3 | 0.186 | | 0.164 | 1.60E-04 |

| | | | | |
|---|---|---|---|---|
| TP63 | | | 1.052 | 1.61E-04 |
| EGR1 | 0.509 | | 1.172 | 1.73E-04 |
| WT1 | | | 1.404 | 1.91E-04 |
| HTT | 0.206 | | 1.671 | 1.94E-04 |
| ZBTB17 | 0.144 | | | 2.19E-04 |
| NFE2L2 | -0.187 | | -0.331 | 2.21E-04 |
| JUN | | | -0.035 | 2.25E-04 |
| PAX3 | | | | 2.36E-04 |
| CREB3L2 | | | | 2.97E-04 |
| ATF6 | -0.146 | | -1.686 | 5.48E-04 |
| BHLHA15 | | | | 5.68E-04 |
| ATF6B | 0.364 | | | 6.42E-04 |
| HIF1A | -0.069 | | 0.093 | 6.51E-04 |
| EPAS1 | 0.186 | | 1.024 | 7.49E-04 |
| NRF1 | -0.054 | | -1.406 | 7.74E-04 |
| YY1 | 0.096 | | | 8.89E-04 |
| NUPR1 | 0.829 | | 1.387 | 9.21E-04 |
| LDB1 | 0.19 | | 0 | 9.37E-04 |
| CREBZF | -0.052 | | | 9.61E-04 |
| LMO2 | -0.026 | | 0 | 9.62E-04 |
| HNF4A | | | 1.399 | 9.88E-04 |
| RRP1B | 0.048 | | | 1.05E-03 |
| SMAD7 | 0.308 | | -1.299 | 1.11E-03 |
| SMARCE1 | -0.223 | | | 1.30E-03 |
| STAT1 | -0.008 | | 0.045 | 1.73E-03 |
| SP3 | -0.205 | | | 2.01E-03 |
| ASCL2 | | | | 2.20E-03 |
| Tcf7 | | | -1.414 | 2.32E-03 |
| KLF11 | 0.441 | | -1.067 | 2.37E-03 |
| E2F1 | 0.199 | | 0 | 2.49E-03 |
| BACH2 | 0.198 | | -0.152 | 2.60E-03 |
| BRCA1 | | | -0.128 | 2.90E-03 |
| NOTCH3 | 0.455 | | | 3.57E-03 |
